# Comprehensive gene heritability estimation reveals the genetic architecture of rare coding variants underlying complex traits

**DOI:** 10.1101/2025.10.07.681018

**Authors:** Zhengtong Liu, Boyang Fu, Moonseong Jeong, Prateek Anand, Aakarsh Anand, Seon-Kyeong Jang, Aditya Gorla, Jiazheng Zhu, Päivi Pajukanta, Pier Francesco Palamara, Noah Zaitlen, Richard Border, Sriram Sankararaman

## Abstract

Whole-exome sequencing (WES) enables high-resolution interrogation of the contribution of rare coding variants to complex trait variation. However, existing methods for heritability estimation attributed to rare-coding variants are often limited by the effects of linkage disequilibrium (LD) and by the sparse nature of rare variant data. We introduce FLEX (*Fast, LD-aware Estimation of eXome-wide and gene-level heritability*), a scalable and flexible framework for estimating and partitioning heritability across genes or sets of genes using WES data. FLEX integrates all coding variants– from common to ultra-rare – within a unified model and corrects for LD-induced effects to improve the accuracy of heritability estimates. In addition, FLEX supports both individual-level and summary statistic data and is computationally efficient for biobank-scale datasets. Through extensive simulations, we show that FLEX is well-calibrated while providing accurate heritability estimates.

We applied FLEX to WES data across *N* = 153, 351 unrelated European ancestry individuals and 20 quantitative traits in the UK Biobank. We identified 64 gene-trait pairs with significant gene-level heritability (*p <* 0.05*/*18, 624 accounting for the number of protein-coding genes tested), among which rare coding variants explained 38% of gene-level heritability, on average. Compared to heritability estimates from genome-wide imputed SNPs, incorporation of rare and ultra-rare coding variants led to a 24.8% increase in heritability on average, while effect sizes at rare and ultra-rare variants are substantially larger (≈ 18x on average). Partitioning across variant effect annotations, we find that predicted loss-of-function variants had stronger individual effects than missense variants (24% on average) while missense variants accounted for a greater share of rare coding heritability. Together, FLEX provides an adaptable and accurate approach for quantifying gene-level heritability, advancing our understanding of the genetic architecture of complex traits, and facilitating the discovery of trait-relevant genes.

## Introduction

The contribution of rare protein-coding variation to complex traits remains incompletely understood. The availability of large-scale whole-exome sequencing (WES) datasets offers the potential to comprehensively characterize the role of rare coding variants and the relative contributions of rare and common coding variants to complex traits [1–3]. A fundamental quantity in understanding the architecture of coding variants is their heritability, *i*.*e*., the proportion of phenotypic variance attributable to this class of variants across the exome (exome-wide heritability or 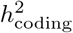). Beyond exome-wide estimates, additional insights into the role of coding variants can be obtained by estimating the variance explained by individual genes (gene-level heritability or 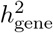). Gene-level heritability estimates offer an approach for discovering trait-relevant genes that is complementary to common-variant genome-wide association studies (GWAS) and gene-based rare-variant association tests (RVATs).

Despite its value, accurately estimating the heritability of coding variants, either at the gene-level or exome-wide, remains challenging. Linkage disequilibrium (LD) among variants within and near gene regions can bias gene-level heritability estimates. Methods such as RARity [4] attempt to mitigate LD-induced inflation via variant pruning, but this can discard causal variants and distort inference. Other approaches rely on posterior sampling of variant effects and LD structure, but are computationally demanding [5]. A further complication arises from the abundance of rare and ultra-rare variants in WES data. Although such variants may have large effects, they are observed in very few individuals, resulting in unstable or inflated heritability estimates [6]. To address this, existing methods often restrict analyses to variants above a minor allele frequency (MAF) threshold (e.g., MAF *>* 0.05) or collapse rare variants into a single burden score [7, 8], assumptions that can limit power or introduce bias.

Here, we introduce **FLEX** (*Fast, LD-aware Estimation of eXome-wide and gene-level heritability*), a scalable and flexible method for estimating and partitioning heritability across individual genes or sets of genes (including all protein-coding genes) using WES data. FLEX integrates all coding variants (common, low-frequency, rare, and ultra-rare) within a unified model and corrects for LD-induced effects to improve the accuracy of heritability estimates. To estimate gene-level heritability 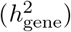, FLEX jointly models variants within each gene and its flanking regions, thereby accounting for local LD and reducing bias. To estimate exome-wide heritability 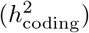, FLEX models variants in more than 18, 000 protein-coding genes together with intergenic variants. FLEX accommodates partitioning of heritability by MAF, LD, or functional genomic annotations, enabling more robust heritability estimates and detailed investigation into genetic architecture. It supports both individual-level genotype data and summary statistics, providing flexibility across data access scenarios. Importantly, FLEX is computationally efficient for biobank-scale datasets.

We first performed extensive simulations to characterize the accuracy, robustness, and computational efficiency of FLEX. We then applied FLEX to the UK Biobank whole-exome sequencing (WES) data to generate estimates of gene-level and exome-wide heritability across diverse quantitative traits. These analyses revealed widespread contributions from rare coding variants, with distinct trait-specific architectures. By providing well-calibrated estimates that account for local LD and leverage variants across the full allele frequency spectrum, FLEX enables more accurate characterization of gene-level contributions to complex traits, facilitating gene prioritization and downstream biological interpretation.

## Results

### Overview of methods

FLEX estimates and partitions heritability across individual genes or sets of genes (including all protein-coding genes) using whole-exome sequencing (WES) data. It employs a linear mixed model to relate phenotypic variation to genetic effects within a protein-coding gene while accounting for contributions from surrounding flanking regions (see “Gene-level linear mixed model” in Methods). The parameters of this mixed model, *i*.*e*. its variance components, correspond to key biological quantities such as the heritability attributable to variants within a given gene (gene-level heritability, 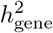) and across all genes (exome-wide heritability, 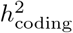). FLEX estimates these variance components via a randomized method-of-moments estimator to enable scalability while retaining accuracy (see “Efficient estimation of variance components” in Methods). Two features distinguish FLEX: it jointly models all coding variants (common, low-frequency, rare, and ultra-rare) within gene regions while also explicitly accounting for LD by modeling variants in flanking intergenic regions. FLEX can be applied to individual-level data (FLEX-h2) or, through an extension we developed, to GWAS summary statistics (FLEX-summ-h2), which is mathematically equivalent to the individual-level implementation (see “Estimating gene-level heritability using summary statistics” in Methods). Finally, to improve power for detecting nonzero 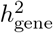, we introduce a variance component test, FLEX-cond-test, which assesses gene-level heritability while controlling for effects from flanking variants.

We applied FLEX to estimate heritability for 18, 624 protein-coding genes across 20 quantitative traits in the UK Biobank with genotypes obtained from *N* = 153, 351 unrelated individuals of European ancestry using a combined dataset of *M* = 21, 512, 331 WES and imputed SNPs. For each protein-coding gene, we defined the gene region to include 10 kb upstream and downstream of the transcription start site (TSS) (see “Gene annotation” in Methods). We stratified variants within each gene region into four MAF bins: common (MAF ≥ 0.05), low-frequency (10^−3^ ≤ MAF *<* 0.05), rare (10^−5^ ≤ MAF *<* 10^−3^), and ultra-rare (MAF *<* 10^−5^) [5, 9]. Ultra-rare variants were collapsed by minor allele count and functional annotation, assuming directional consistency of effects within each group, following prior approaches [6, 8] (see “Collapsing ultrarare variants into pseudo-markers” in Methods).

For each trait, we used FLEX to estimate gene-level heritability by fitting variance components corresponding to the MAF bins, allowing both bin-specific and overall 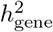 estimates. We then extended this framework to exome-wide analyses, estimating 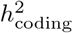 from all protein-coding variants together with intergenic variants captured in the imputed data. Finally, we partitioned heritability across genomic annotations such as MAF and LD, or variant annotations – protein loss-of-function (pLoF), missense, and synonymous – to dissect the relative contributions of different variant classes to trait heritability.

### Accurate, robust, and efficient gene-level heritability estimation in simulations

To evaluate the statistical calibration of FLEX, we performed chromosome-level simulations where we simulated phenotypes under a null model of no gene-level heritability using merged WES and imputed genotypes from chromosome 21 (see Section “Genotype and phenotype data” in Methods). In these simulations, we varied the background heritability (the heritability attributed to variants that lie outside these genes) while exploring a range of genetic architectures that varied in the dependence of allelic effects on MAF and LD and the proportion of causal variants (see Section “Simulation details: Calibration” in Methods). FLEX-h2 produced well-calibrated estimates, with mean heritability centered near zero and false positive rates (FPRs) aligned with the nominal 5% level (*p* = 0.30–0.89 for one-sided test of FPR *>* 0.05; Figure 1a; Figure S1). The summary-based extension FLEX-summ-h2 yielded similarly unbiased estimates (Figure S2a) while the conditional test FLEX-cond-test maintained controlled FPR across all backgrounds (Figure S4). We further verified that the tests of 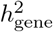 from FLEX-h2 and FLEX-cond-test were calibrated across a wide range of *p*-values, although we note that the Wald test in FLEX-h2 yielded *p*-values that tend to be slightly conservative (Figure S3).

**Figure 1:**
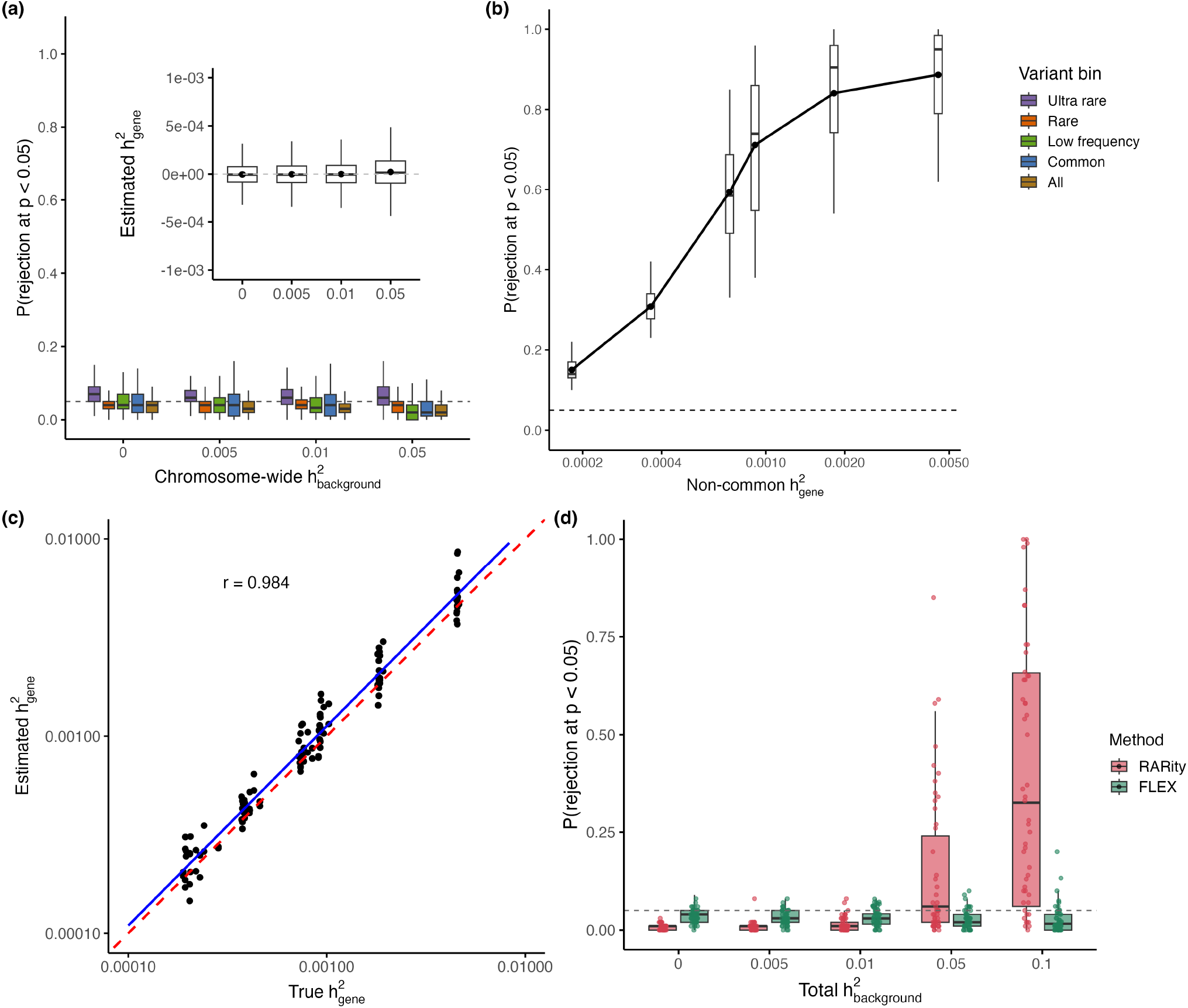
Calibration and power of FLEX in simulations. **(a)** False positive rates (FPR) under varying levels of background heritability. FPR was evaluated across 100 replicates per setting, with 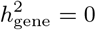. Each box summarizes the proportion of rejections at *p<* 0.05 across 117 genes on chromosome 21 and various genetic architectures. Variant sets are stratified by MAF; “All” includes all variants within each gene. The horizontal dashed line marks the nominal 5% threshold. The inset displays the distribution of point estimates, with dots indicating the mean. **(b)** Power to detect 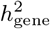 as a function of non-common 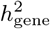. The *x*-axis denotes the simulated non-common 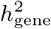 for causal genes, defined as the heritability attributable to variants with MAF *<* 0.05 within gene regions. Boxplots show results across 100 replicates per setting under different genetic architectures, with the dashed line indicating the 5% significance threshold. **(c)** Accuracy of 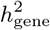 estimates plotted against the true simulated values across all genes and replicates (*r* = 0.984). A best-flt regression line (blue) and the identity line (dashed red) are shown. **(d)** Comparison of false positive rates between FLEX and RARity with increasing levels of background heritability. Each box shows the proportion of rejections at *p<* 0.05 across 100 replicates.

We next assessed the power of FLEX to detect gene-level heritability (see Section “Simulation details: Power and accuracy” in Methods). Across a range of simulated heritability values, FLEX-h2 produced point estimates that closely tracked the true values (Figure 1c; Figure S2b) and achieved adequate power to detect genes with non-common 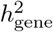 as low as 1 *×* 10^−3^ at a *p*-value threshold of *p <* 0.05 (Figure 1b) while the conditional test (FLEX-cond-test) demonstrated additional increase in power over the Wald test (Figures S5 and S6).

Importantly, we note that FLEX is accurate even though the model used for simulation (which assumes effect sizes vary as a continuous function of MAF and depend on local LD) differed from the model used for estimation (that uses four MAF bins to model how effect sizes vary with MAF), justifying the use of the four MAF bins in our analyses. To further evaluate our modeling choices, we tested the impact of the number of random vectors used in the trace estimator and the size of the LD flanking region (see “Simulation details: Sensitivity to modeling parameters” in Methods). Across simulations with heritability from both common and non-common variants (*h*^2^ = 0.01 each), using as few as 10 random vectors yielded highly concordant point estimates compared to 100 or 500 vectors, with only a modest increase in standard error (≈ 10%; Figure S8). We also evaluated flanking regions of 10 kb, 100 kb, and 1 Mb under a null model with background heritability in intergenic regions. While all window sizes achieved average FPRs near the nominal threshold of 0.05, under high values of background heritability, the variability of the FPRs was stable for window sizes larger than 100 kb, motivating the use of 100 kb as the default window size in our analyses (Figure S9). We further assessed the impact of collapsing ultra-rare variants (see “Simulation details: Effect of ultra-rare variant collapsing” in Methods) and observed a modest decrease in power due to collapsing (Figure S7).

Finally, we benchmarked the computational scalability of FLEX-h2 on individual-level data, noting that the summary-statistics implementation, FLEX-summ-h2, is even faster (see “Simulation details: Runtime benchmarks” in Methods). Across sample sizes of 30,000 to 100,000 individuals, the runtime per gene ranges from 5 to 25 seconds (Figure S10a). FLEX-h2 also scales efficiently with the number of SNPs per gene region; for regions containing up to 6,000 variants, the runtime remained around 30 seconds at a sample size of 100,000 (Figure S10b).

### Comparison to existing methods in simulations

Building on the simulation results above, we next benchmarked the performance of FLEX against two recent approaches for modeling rare variant contributions in whole-exome sequencing studies. BHR aggregates rare variants within gene regions into a burden score to estimate burden heritability (which is a subset of gene-level heritability), while RARity excludes singletons and applies LD pruning to limit the impact of LD [9, 4].

We first compared the calibration of these methods under null simulations. In the presence of modest background polygenicity 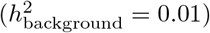, both BHR and RARity produced calibrated estimates for rare and ultra-rare variant bins but exhibited inflated false positive rates for low-frequency and common variants, consistent with BHR’s design (Figure S11). In contrast, FLEX remained well-calibrated across all MAF bins. As background heritability increased, RARity’s FPR became progressively inflated, while FLEX maintained robust calibration (Figure 1d). We excluded BHR from these comparisons at higher 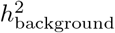settings, as it does not model common variant architectures.

We next compared accuracy across methods in the rare and ultra-rare MAF bins, where all approaches were well-calibrated (see “Simulation details: Benchmarking calibration and power against existing methods” in Methods). Across simulation settings, FLEX consistently recovered more gene-level heritability than the next best method: the point estimates captured 91.0% (86.5%) of the simulated heritability in the rare (ultra-rare) bin on average across simulations, compared to 89.4% (60.5%) for RARity (Figure S12).

### Gene-level heritability estimation in UK Biobank

We applied FLEX to estimate gene-level heritability for 18, 624 protein-coding genes across 20 quantitative traits in the UK Biobank (see “Genotype and phenotype data” and “Gene annotation” in Methods). For each gene, we computed heritability stratified by MAF to estimate gene-level heritability for common, low-frequency, rare, and ultra-rare variants, respectively.

We first evaluated the impact of accurately modeling LD in real data by comparing estimates obtained from FLEX with and without flanking-region adjustment. Specifically, the unconditional model applied FLEX using variants restricted to the gene body while the conditional model included the flanking regions to account for LD (see Section “Gene-level linear mixed model” in Methods). LD conditioning led to consistently lower estimates of 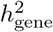across gene-trait pairs (paired *t*-test, *p* = 1.90 *×* 10^−13^; Figure 2a, Figure S13) and to a 24.3% reduction in the number of nominally significant gene-trait associations (*p <* 0.05). For instance, inflation in − log_10_ p-values for C-reactive protein was markedly attenuated following LD correction (Figure 2b). Despite this reduction, core signals were retained: across all 20 traits, 79.0% of marginally significant genes identified after LD correction overlapped with those identified without conditioning (Figure S14). Stratifying our results by MAF bin, we find significantly lower estimates following LD correction for all MAF bins except ultra-rare variants (Figure S13), which were minimally influenced by LD due to their very low allele counts.

**Figure 2:**
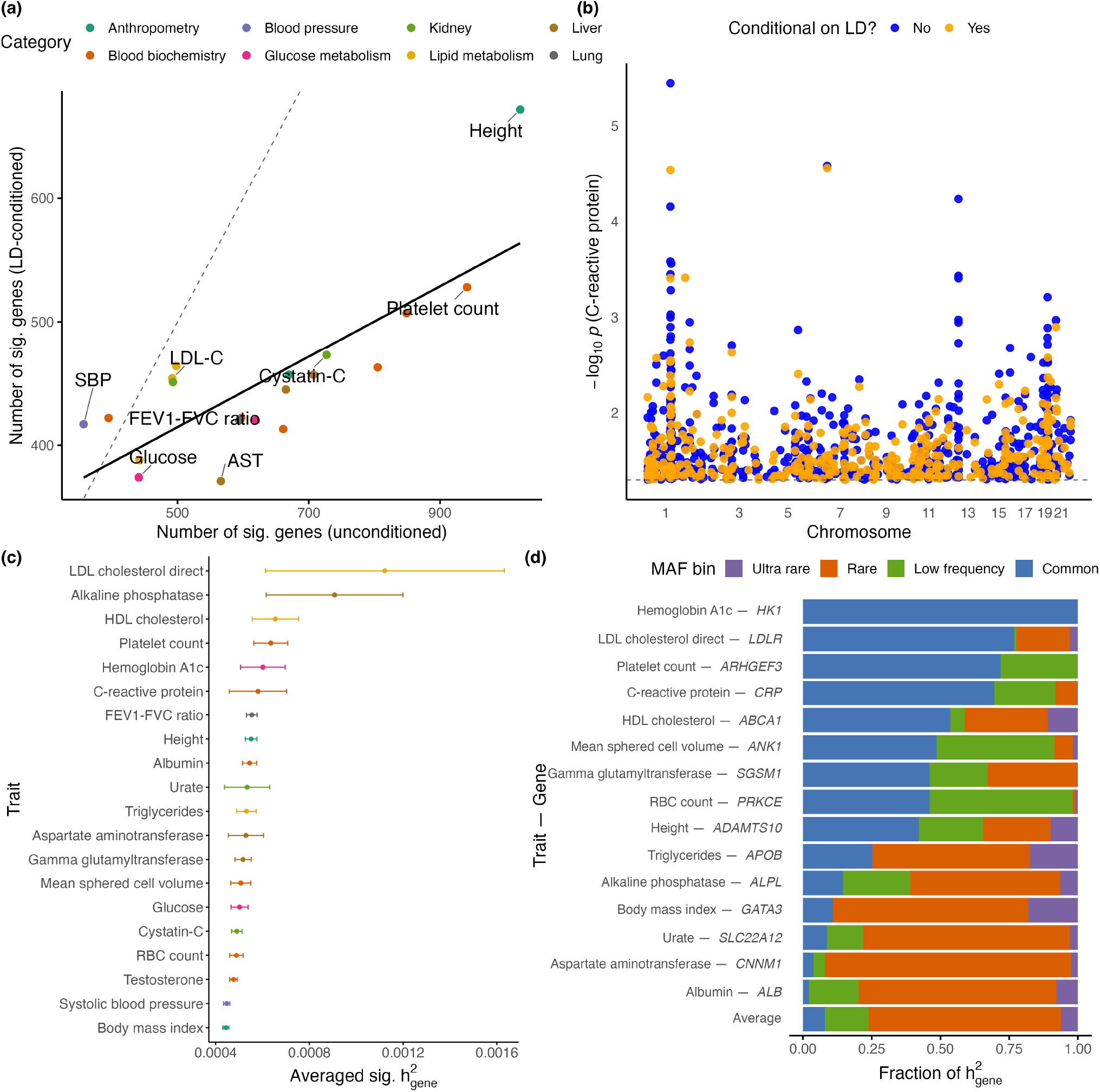
Trait-level patterns of gene-level heritability and the impact of LD conditioning. **(a)** Number of significant genes (*p<* 0.05) identified by FLEX-h2 in the unconditioned versus LD-conditioned analysis. Each point represents a trait and is colored by trait category. The dotted diagonal line denotes equality (*y* = *x*), and the solid line indicates the best-flt linear regression across traits. **(b)** Conditional versus unconditioned *p*-values for gene-level heritability of C-reactive protein. Each point represents a gene, with blue indicating the unconditioned model and orange indicating the LD-conditioned model. **(c)** Average 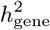 across genes with significant 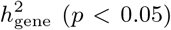 per trait. Traits are sorted by average 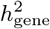 among significant genes. Error bars denote 95% confidence intervals. **(d)** Fraction of 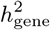attributable to each minor allele frequency (MAF) bin for selected trait-gene pairs. Each horizontal bar represents a single trait-gene pair with significant 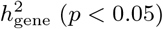, and colors indicate the contribution from each MAF bin. The bottom bar shows the average across all significant trait-gene pairs (*p<* 0.05).

All subsequent analyses were performed using LD-conditioned estimates of 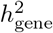. Across all gene-trait pairs, 2.4% (9,012 pairs) exhibited nonzero 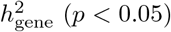, with an average heritability estimate of 5.81 *×* 10^−4^ (Figure 2c). The mean 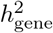 ranged from 4.44 *×* 10^−4^ for body mass index (BMI) to 1.12 *×* 10^−3^ for LDL cholesterol direct (LDL-C). Partitioning 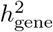 across MAF bins, we find that rare variants make a substantial contribution to gene-level heritability: among genes with nominally significant 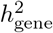 estimates, rare and ultra-rare variants explained 76.1% of 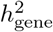 on average. However, the relative contributions of rare and common variants varied widely across gene-trait pairs: *HK1*-HbA1c was almost entirely driven by common variants whereas *CNNM1*-aspartate aminotransferase (AST) showed strong enrichment in rare variants, which explained 89.5% of its 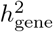 (Figure 2d).

### Quantifying trait polygenicity through gene-level heritability

Beyond identifying individual gene-trait associations, we sought to understand polygenicity at the gene level, *i*.*e*., whether heritability is concentrated in a few genes or distributed across many genes. We observed several examples of genes that harbor substantial heritability for specific traits. For instance, *ALPL*, a key regulator of alkaline phosphatase (ALP) activity, accounted for a substantial fraction (13.1%) of the total 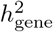 of ALP. Similarly, *APOE* explained 28.1% of the total 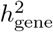 for LDL-C, while *CRP* explained 7.0% for C-reactive protein levels. To systematically quantify gene-level polygenicity for a specific trait, we ranked genes with nominally significant 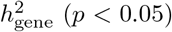 and computed *τ*_80_: the minimum number of genes required to explain a substantial fraction (80%) of the cumulative gene-level heritability. Across traits, we find *τ*_80_ = 295 so that 295 genes explain a substantial fraction of the cumulative gene-level heritability with lipid traits being the least polygenic (average *τ*_80_ = 247 genes) while anthropometric traits were the most polygenic (average *τ*_80_ = 385 genes) (Figure 3c). The relative rankings across traits were largely preserved across alternative cumulative heritability thresholds (e.g., 50%) and are broadly consistent with prior variant-level analyses [10, 11]. We find that polygenicity structure was also preserved for rare and ultra-rare coding variants (MAF *<* 10^−3^): *τ*_80_ attributed to these variants was highly correlated with that from all variants (Pearson’s *r* = 0.86, *p* = 1.30 *×* 10^−6^; Figures S15 and S16). These analyses reveal susbtantial polygenicity for complex traits at the gene-level both when considering all variant classes and while restricting to rare variants.

**Figure 3:**
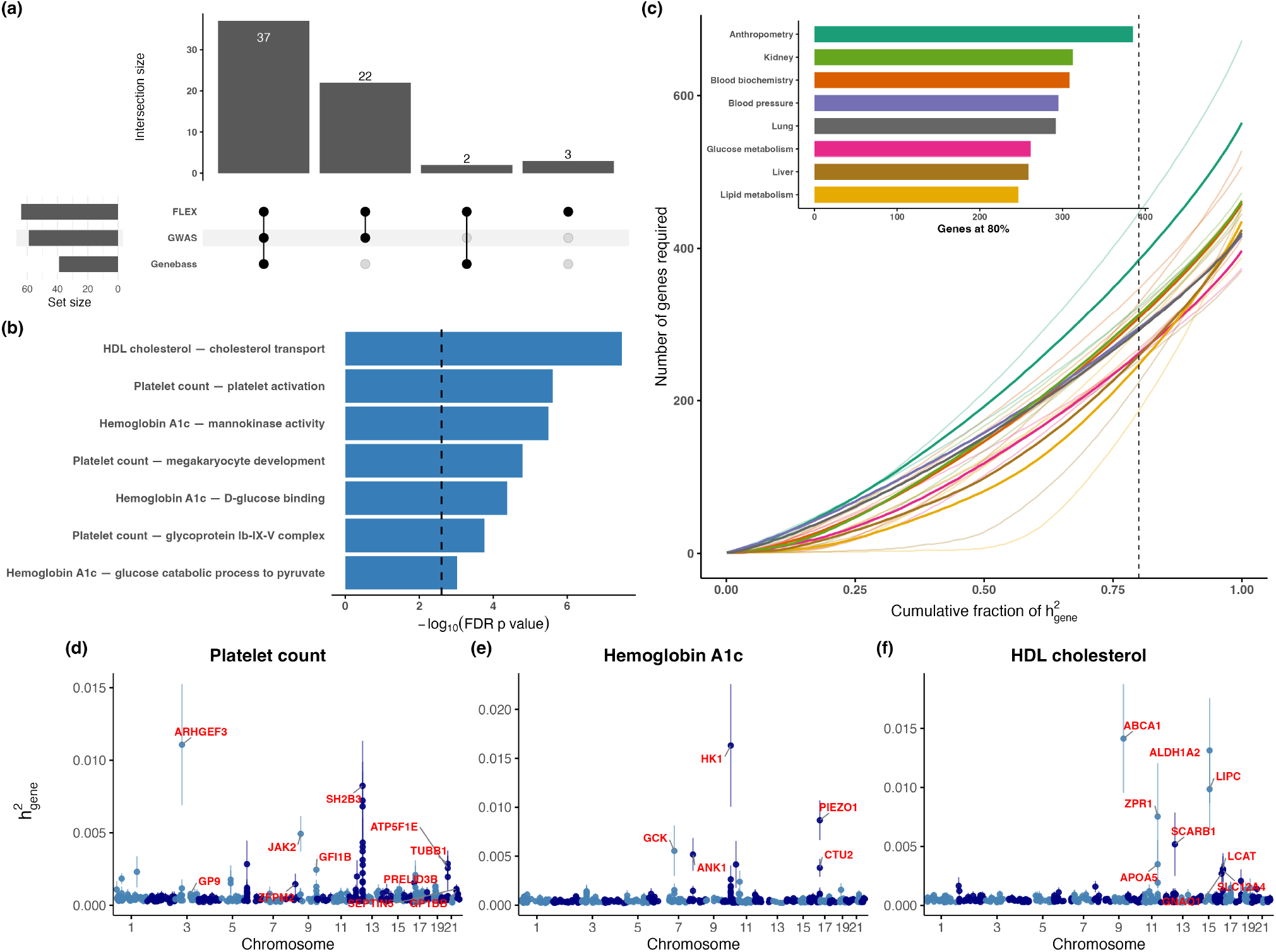
Validation, functional interpretation, and polygenic structure of gene-level heritability. **(a)** Overlap of genes with significance across FLEX (*p<* 0.05*/*18, 624), GWAS (*p<* 5 *×* 10^−8^), and Genebass gene-based tests (*p<* 2.5 *×* 10^−7^). The top panel shows the number of genes in each intersection, and the bottom panel shows the total number of significant genes per method. Only genes passing their respective significance thresholds for a given trait were included in the comparison. **(b)** Selected Gene Ontology (GO) terms significantly enriched among genes with genome-wide significant heritability (*p<* 0.05*/*18, 624) for a given trait. Bar length represents the − log_10_(FDR) of enrichment, and the horizontal dotted line marks the significance threshold after correcting for the number of traits (FDR *p <* 0.05*/*20). Enrichment was performed using only genome-wide significant genes identified for that trait. **(c)** Cumulative fraction of 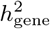 explained by top-ranked genes. Each faint line represents an individual trait, and solid lines indicate the average curve for each trait category, colored accordingly. The inset shows the number of genes required to explain 80% of 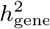 per category, summarizing polygenicity across traits. **(d-f)** Genome-wide Manhattan-style plots for gene-level heritability estimates of three representative traits. Each point represents a gene, with vertical lines indicating standard errors of the heritability estimates. Genes with genome-wide significant heritability (*p<* 0.05*/*18, 624) are labeled in red.

### Comparing gene-level heritability with GWAS and gene-based association tests

Common-variant GWAS and gene-based rare-variant association tests (RVAT) are standard approaches for identifying gene-trait associations. These methods typically detect variants or genes with statistically significant effects, prioritizing loci with large or annotation-enriched effect sizes. In contrast, heritability estimation aggregates the contribution of all variants within a gene, regardless of their individual significance. Thus, gene-level heritability offers a complementary perspective on genetic architecture: it can reveal polygenic contributions that may be missed by association-based approaches, and its signals can be supported or contextualized by existing statistical associations.

To explore this complementarity, we examined the extent to which genes with genome-wide significant 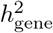 were supported by existing GWAS and gene-based RVAT tests. Across all traits, we first identified 64 gene-trait pairs with genome-wide significant 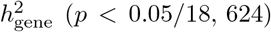. We observe that estimates of 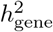 span two-orders of magnitude: from *GATA3*-BMI 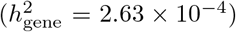 to *APOC1*-LDL-C 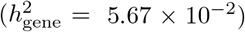 (Figure S17) with rare and ultra-rare variants contributing about 38% of 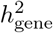 on average. Among these gene-trait pairs, 59 overlapped with genome-wide significant variants reported for the same traits in the GWAS Catalog [12] (Figure 3a; see “Overlap analysis” in Methods). Among these overlapping pairs, we observed a positive correlation between 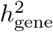 estimates and the number of GWAS hits per gene (*r* = 0.48, *p* = 3.21 *×* 10^−5^; Figure S18), suggesting that strong marginal associations contribute to increased gene-level heritability.

We further compared the significant gene-trait pairs with gene-based RVAT results from Genebass for pLoF, missense, and synonymous variants [13] (see “Overlap analysis” in Methods). We observed overlap with 21, 34, and 17 gene-trait pairs for pLoF, missense, and synonymous SKAT-O tests, respectively (Figure S19, *p <* 2.7 *×* 10^−7^). Since SKAT-O *p*-values reflect both effect size and uncertainty [14], we correlated gene-level *z*-scores 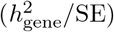 to SKAT-O *p*-values to find significant correlations for pLoF (*r* = 0.69, *p* = 4.79 *×* 10^−4^) and missense (*r* = 0.44, *p* = 1.40 *×* 10^−2^) variants, but not for synonymous variants (*p* = 0.13) (Figure S20). Three gene-trait pairs displayed genome-wide significant 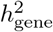 estimates but did not have significant associations in GWAS and RVAT (Figure 2a): *ZFYVE27*-mean sphered cell volume (MSCV) 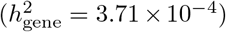, *TRAPPC6A*-LDL-C 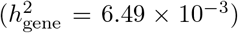, and *GATA3*-BMI 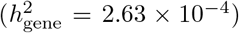. *GATA3*-BMI, while showing no association in GWAS or RVATs, has been implicated in adipocyte differentiation and metabolic regulation [15] with a variant within this gene (rs12412241) associated with BMI at a sub-genome-wide significance level (*p* = 3 *×* 10^−6^) in the GWAS Catalog. These findings underscore the value of gene-level heritability as a complementary tool to detect genes with distributed effects that may be overlooked by GWAS or RVATs alone.

### Functional enrichment of gene sets with significant gene-level heritability

To better understand the biological relevance of genes identified in our heritability scan, we performed Gene Ontology (GO) enrichment analysis on genes with genome-wide significant heritability (*p<* 0.05*/*18, 624) to identify overrepresented molecular functions and biological processes. To characterize trait-specific biological pathways, we applied `g:Profiler` to significant 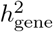 genes, incorporating GO, KEGG, and TRANSFAC annotations [16] (see “Gene set enrichment” in Methods). For each trait, we reported FDR-corrected *p*-values and applied a multiple testing threshold of *p<* 0.05*/*20, correcting for the number of traits analyzed. This analysis revealed consistent enrichment of biologically relevant processes. For HbA1c, enriched terms included mannokinase activity, D-glucose binding, and the glucose catabolic process to pyruvate, highlighting key steps in cellular glucose metabolism [17] (Figures 3b,e). Blood platelet traits showed enrichment in platelet activation, megakaryocyte development, and the glycoprotein Ib-IX-V complex pathway, consistent with the central role of this receptor complex in platelet adhesion and thrombus formation [18, 19] (Figures 3b,d). For HDL cholesterol, significant genes were enriched for cholesterol transport, reflecting their roles in lipoprotein remodeling and cholesterol homeostasis [20] (Figures 3b,f). These results demonstrate that gene-level heritability estimates can recover trait-relevant biology and offer mechanistic insights into disease-associated pathways.

### Contribution of rare coding variants to genome-wide heritability

To systematically quantify the overall contribution of rare coding variants, we applied FLEX across the genome where we jointly analyzed all SNPs that were represented in the WES data across protein-coding genes along with imputed SNPs in intergenic regions (effectively analyzing the set of all protein-coding genes). Specifically, we concatenated genotypes spanning all protein-coding genes and the intervening intergenic regions – retaining only imputed SNPs with MAF *>* 10^−5^ in the latter – and merged overlapping intervals across adjacent loci to define unified coding regions. To allow for SNP effect sizes that vary with MAF, and LD, we followed prior studies [21, 22] and partitioned SNPs into four MAF bins (common, low-frequency, rare, and ultra-rare) and further subdivided into high- and low-LD strata using local LD score thresholds. We also adjusted estimates of height to account for assortative mating assuming that the same adjustments apply across all bins [23].

Across the traits analyzed, the inclusion of rare and ultra-rare coding variants led to a 24.8% increase in heritability estimates beyond that captured by common and low-frequency variants (MAF ≥ 10^−3^) with traits such as MSCV showing a substantial 61.5% increase (Figure 4a). To examine differences in effect distribution, we estimated the average squared per-allele effect sizes *β*^2^ for each MAF bin (see “Estimation of exome-wide heritability in the UK Biobank” in Methods). Rare and ultra-rare coding variants 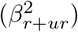 exhibited substantially larger effects compared to common and low-frequency coding variants 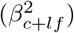 (Figure 4b, meta-analyzed 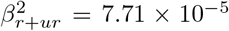 for rare and ultra-rare vs 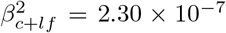 for common and low-frequency variants), in line with prior reports of larger effects among rare variants subject to negative selection [24, 25, 9].

**Figure 4:**
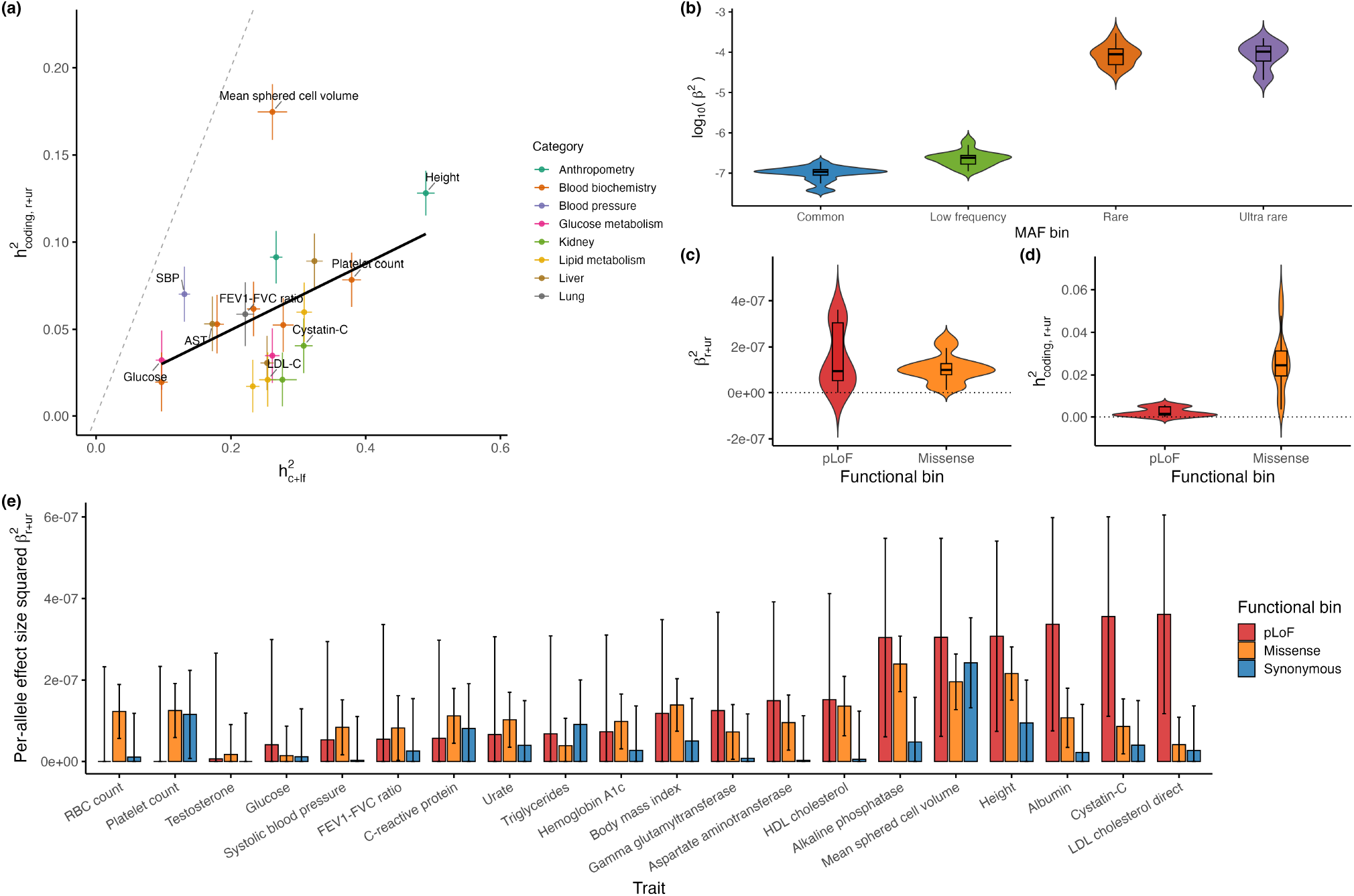
Patterns of exome-wide heritability across traits, allele frequency bins, and functional annotations. **(a)** Heritability attributed to rare and ultra-rare coding variants across the exome 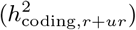 compared to common and low-frequency variants across the genome 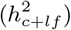. Each point represents a trait, with error bars indicating 95% confidence intervals. The dotted diagonal line denotes equality (*y* = *x*), and the solid line indicates the best-flt linear regression across traits. This comparison highlights the exome-wide contribution of rare coding variation. **(b)** Distribution of squared per-allele effect sizes of coding variants across the exome (*β*^2^) stratified by MAF bin. **(c)** Distribution of squared per-allele effect sizes 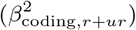 and **(d)** heritability estimates 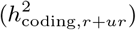 for pLoF and missense, rare and ultra-rare variants. **(e)** Squared per-allelic effect size among rare and ultra-rare coding variants, partitioned by functional consequence: pLoF, missense, and synonymous.

Although the squared effect sizes 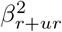 and 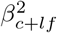 were positively correlated across traits (*r* = 0.50, *p* = 2.40 *×* 10^−2^), the relative contribution of rare versus common variants varied substantially by trait category. To quantify this variation, we computed the ratio 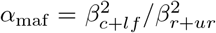 as a summary measure of relative magnitudes of the effects of common vs rare variants. Kidney traits reflected a greater contribution from common coding variants (meta-analyzed *α*_maf_ = 4.39 *×* 10^−3^) while blood biochemistry traits showed the lowest (meta-analyzed *α*_maf_ = 2.36 *×* 10^−3^), indicating stronger effects from rare coding variants in these traits (Figure S21) [26]. We did not observe substantially larger effects at ultra-rare vs rare coding variants (meta-analyzed 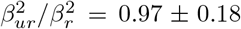), consistent with the under-estimation of ultra-rare coding variant heritability that we see in our simulations and likely due to the collapsing of these variants.

### Rare coding heritability across functional annotations

We next examined how functional annotations influence the contribution of rare coding variants to heritability. Variants were jointly stratified by predicted molecular consequence – pLoF, missense, synonymous – and by allele frequency: common/low-frequency (combined), rare, and ultra-rare (see “Variant annotation” in Methods). This partitioning enabled us to assess how both variant frequency and function shape heritability patterns. We defined 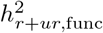 as the proportion of trait variance explained by rare and ultra-rare variants in a given functional class.

We estimated the average squared per-allelic effect size for the three functional annotations (pLoF, missense, and synonymous) combining estimates across 20 traits by meta-analysis (see “Estimation of exome-wide heritability in the UK Biobank” in Methods). Consistent with prior findings [9], rare pLoF variants showed larger average effect sizes than missense variants (Figure 4c, meta-analyzed squared per-allelic effect size 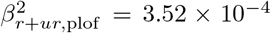 for rare and ultra-rare pLoF variants compared to 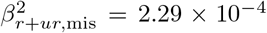 for missense variants). However, missense variants explained a larger proportion of heritability among rare coding variants (meta-analyzed 76.3% for rare missense and 6.1% for rare pLoF variants; Figure 4d, Figure S22).

We further explored trait-specific differences in the contribution of rare functional variants, focusing on pLoF and missense categories (Figure 4e). We computed the ratio 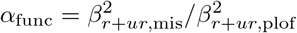 as a measure of relative magnitudes of the effects of rare, missense vs pLoF variants (restricting to traits with positive estimates of 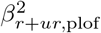 and 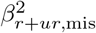). LDL-C exhibited the strongest relative enrichment for pLoF effects (*α*_func_ = 0.05), consistent with known roles of loss-of-function mutations in lipid genes such as *PCSK9, LDLR*, and *APOB* [27–29]. In contrast, three traits – systolic blood pressure, C-reactive protein, and serum urate – exhibited *α*_func_ *>* 1, indicating stronger effects from rare missense variants than from pLoFs (Figure S23). For serum urate, this pattern is consistent with external evidence: two rare, independent missense variants in *SLC22A12* – also the top urate-associated gene in our analysis 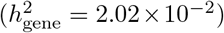 – showed exome-wide significant associations with large effects, one of which has been linked to hypouricemia [30, 31]. Together, these findings provide a systematic comparison of the relative contributions of rare coding variants across functional classes.

## Discussion

We have presented FLEX, a scalable and versatile framework for estimating and partitioning heritability at the level of individual genes or predefined gene sets using whole-exome sequencing data. A key feature of FLEX is its ability to jointly model all coding variants – ranging from common to ultra-rare – within gene regions. To incorporate ultra-rare variants, which are typically sparsely observed, we adopt a pseudo-marker strategy that collapses variants by minor allele count and functional annotation under an assumption of directional consistency. This approach aims to improve power while limiting potential bias, adapting ideas from burden testing into a heritability framework. Another component of FLEX is the explicit adjustment for LD, achieved by modeling variants in nearby intergenic regions. We show that this adjustment controls for false positives and improves the accuracy of our estimates of 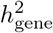. In our analyses, we found that short genes and those with high-LD variants were particularly susceptible to inflation, affecting both common and rare variants (Figures S24 and S25, see Supplemental Notes, “Impact of linkage disequilibrium on gene-level heritability estimation”). These findings suggest that flanking-region LD adjustment plays an important role in controlling false positives and improving interpretability for gene-level analyses.

When applied to the UK Biobank WES data, FLEX identified multiple gene-trait pairs with significant gene-level heritability. While gene-trait pairs with significant 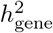 were generally also implicated in GWAS and RVATs, our finding of three gene-trait pairs with significant 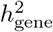 that are missed by current GWAS and RVATs suggests that gene-level heritability estimates can serve as a complementary tool for discovering trait-relevant genes. Although previously identified as a crucial gene in lipid metabolism and cardiometabolic risk [32], our finding of high 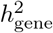 at *APOC1* points to the relevance of its genetic effects for LDL-C, offering additional support for its potential as a therapeutic target. On the other hand, our finding of substantial gene-level polygenicity, with most of the gene-level heritability distributed across hundreds of genes, suggests that larger sample sizes will be needed to identify trait-relevant genes.

By stratifying heritability estimates by MAF, we observed that rare variants – despite their modest constribution to genome-wide heritability [25] – can contribute substantially to gene-level heritability, on average, and to the heritability attributed to specific gene-trait pairs. Notably, the effect sizes at rare and ultra-rare variants tend to be substantially larger than at common and low-frequency variants, highlighting the disproportionate influence of rare coding variants. Among variant classes, missense variants accounted for the largest share of total 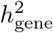, while pLoF variants, although fewer, exhibited the strongest per-allele effects. In contrast, synonymous variants appeared to contribute little. These observations offer additional insight into how different types of coding variation shape phenotypic variability.

Despite its utility, FLEX has several methodological limitations that warrant further development. First, the current implementation ignores individuals with missing phenotypes. Incorporating phenotype imputation strategies could increase the effective sample size and thereby improve power [33–35]. Second, while FLEX can accommodate binary traits and achieves good calibration and power under low case-prevalence conditions (prevalence = 0.1, Figure S26), its performance may degrade for extremely unbalanced case– control settings. Developing extensions to better address such imbalance [36, 8] would improve the method’s relevance for rare disease gene discovery. Finally, the scalability of FLEX relies on a randomized method-of-moments estimator, which is computationally efficient but statistically less efficient than restricted maximum likelihood (REML) under ideal assumptions [37]. Designing likelihood-based estimators that remain feasible at biobank scale is an important future direction.

Our current applications also highlight several directions for future work. The study focuses on the role of rare coding variants. The growing availability of whole-genome sequencing (WGS) data offers the opportunity to understand the role of rare non-coding variants in complex traits. While FLEX is compatible with WGS-based rare variants, additional work is needed to incorporate broader functional genomic annotations such as noncoding and regulatory elements. Furthermore, our analyses were restricted to individuals of European ancestry. Applying FLEX to whole-exome and whole-genome sequencing datasets from diverse ancestries [38] will be essential for comprehensively characterizing shared and population-specific aspects of complex trait genetic architecture.

## Methods

### Gene-level linear mixed model

We consider the following linear mixed model linking genotypes within and around a gene to the phenotype:

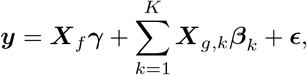

where ***y*** ∈ ℝ ^*N*^ denotes standardized phenotypes for *N* individuals. The matrices 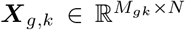 and 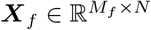 contain standardized genotypes for the *k*-th functional annotation within the gene and for the flanking region, respectively. SNPs in the gene region are partitioned into *K* annotations, and the flanking region is defined as a fixed window upstream and downstream of the gene. The vectors ***γ*** and ***β***_*k*_ (of lengths *M*_*f*_ and *M*_*gk*_, respectively) represent SNP effect sizes in the flanking region and in the *k*-th annotation of the gene region. The residual term ***E*** captures noise. We assume 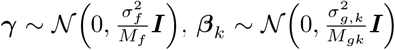 and 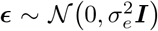, where the variance components 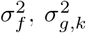, and 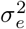 correspond to the contributions from the flanking region, the *k*-th gene annotation, and residual noise, respectively.

To estimate the gene-level heritability while also partitioning the heritability across MAF bins, we split each gene region into common, low-frequency, rare, and ultra-rare variants defined as SNPs with MAF ≥ 0.05, ≥ 1*e*^−3^, ≥ 1*e*^−5^ and *>* 0, respectively [5, 9]. Ultra-rare variants corresponded to variants with minor allele count (MAC) ∈ {1, 2, 3} in our dataset. We collapsed ultra-rare variants into pseudo-markers, as described in the section “Collapsing ultra-rare variants into pseudo-markers”. Let ***X***_*g,c*_, ***X***_*g,lf*_, ***X***_*g,r*_, ***X***_*g,ur*_ denote the genotypes of common, low-frequency, rare and ultra-rare variants in gene *g*, with 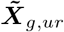 denoting the collapsed genotypes for ultra-rare SNPs. Thus, we fit the following gene-level linear mixed model:

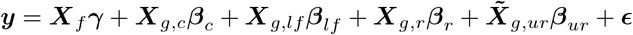

In this model, the variance components 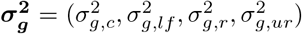 corresponding to ***β***_***c***_, ***β***_***lf***_, ***β***_***r***_ and ***β***_***ur***_ are the key parameters of interest. For each MAF bin, we can then compute the gene-level heritability associated with each MAF bin as: 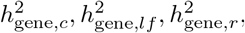 or 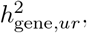, respectively. The overall gene-level heritability estimate, 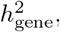, is obtained by adding up the heritability estimates associated with each MAF bin.

### Collapsing ultra-rare variants into pseudo-markers

We collapsed ultra-rare variants – those with MAF *<* 10^−5^ or MAC *<* 4 in our dataset – into pseudo-variant markers [6, 9, 8]. For each gene, ultra-rare variants were grouped based on their functional annotation (e.g., missense, loss-of-function) and MAC bin. Variants within the same group were collapsed by recording, for each individual, the maximum allelic count observed across all variants in that group, consistent with prior works [8]. Formally, let *𝒱* _*g*_ denote the set of ultra-rare variants in gene *g* that fall into the same annotation category. For individual *i*, the collapsed genotype 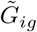 is computed as:

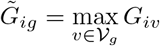

where *G*_*iv*_ is the allelic count (0, 1, or 2) for variant *v* in individual *i*. To improve statistical stability, annotation groups with fewer than ten variants were further aggregated into a single pseudo-marker prior to collapsing.

### Efficient estimation of variance components

We use the methods-of-moments (MoM) to estimate the variance components in the linear mixed model [39–41]. Assume the phenotype vector ***y*** is standardized such that E[***y***] = 0 and Var[***y***] = 1. The MoM estimation aims to match the population variance according to the gene-level linear mixed model 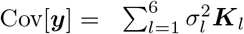 with the empirical covariance estimated as ***yy*** ^⊤^. We denote the variance components by 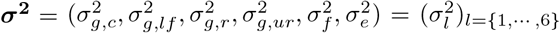 and the GRM matrix of the *l*-th variance component by ***K***_*l*_ (the GRM matrix for the noise component is ***I***_*N*_). The MoM parameter estimates are obtained by minimizing the squared Frobenius norm between the population and empirical variance.

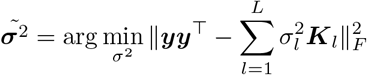

The optimization problem can then be reduced to solving the normal equations

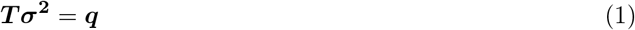

with ***T*** _*ij*_ = tr [***K***_*i*_***K***_*j*_] and ***q***_*p*_ = ***y*** ^⊤^ ***K***_*p*_***y***, for *i, j, p* ∈ {1, ···, 6}. The heritability of the *l*-th partition of genotypes is then defined as

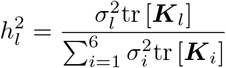

Naively computing the trace ***T***_*ij*_ = tr [***K***_*i*_ ***K***_*j*_]requires constructing the GRM ***K*** and multiplying two GRMs, which incurs a cost of *𝒪* (*N* ^3^) and constitutes the main computational bottleneck of the algorithm (forming ***X***^**⊤**^ ***X*** costs *𝒪* (*N* ^2^*M*) and ***KK*** costs *𝒪* (*N* ^3^), where *N* and *M* denote the numbers of individuals and variants, respectively). Therefore, we implemented an efficient computation of the trace ***T*** _*ij*_ = tr [***K***_*i*_***K***_*j*_] using the randomized trace estimator [42]. Assume ***z***_*k*_ is a N-vector such that 𝔼 [***z***_*k*_] = **0** and 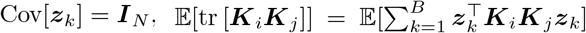, or 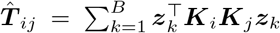 is an unbiased estimator of ***T*** _*ij*_. In practice, we sampled *B* random vectors ***z***_*k*_ from 𝒩 (**0, *I***_*N*_). This reduced the runtime computation of ***T*** _*ij*_ from *𝒪* (*N* ^3^) to *𝒪* (*NMB*). Furthermore, we utilized the specific structure of the genotype matrix, which contains only values of 0, 1, or 2, by applying the Mailman algorithm [43]. This further reduced the time complexity of matrix-vector multiplication per iteration from 𝒪 (*NM*) to 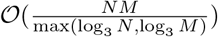. The overall runtime complexity for our algorithm is then 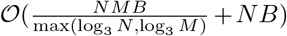for a single trait-gene pair, a significant improvement from the direct computation that costs *O*(*N* ^3^). We refer to the gene-level heritability estimator with accompanying standard errors as **FLEX-h2** throughout our analysis.

### Standard errors of the estimates

We compute asymptotic standard errors (SEs) for the variance component estimates [44]. For the variance component estimates of a single trait, 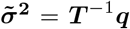 so that 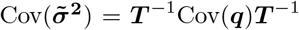. We then compute the *ij*-th entry of the covariance matrix of ***q*** using the variance of the quadratic form:

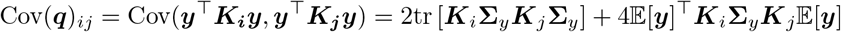

where **Σ**_*y*_ = Cov[***y***]. Since 𝔼 [***y***] = **0** and by the cyclic property of the trace operator, Cov[***q***]_*ij*_ = 2tr [**Σ**_*y*_***K***_*i*_**Σ**_*y*_***K***_*j*_]. Then we use ***yy***^**⊤**^ as the empirical estimate of **Σ**_*y*_. Under the gene-level linear mixed model, we use 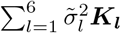 as the approximation of ***yy***^**⊤**^ where 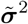 is the MoM estimator. Therefore, we estimate the Cov[***q***] as

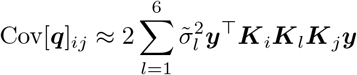

Because ***T*** is fixed across traits and very small, these SE computations add negligible overhead once ***T*** and the necessary matrix–vector products have been precomputed.

### Efficient variance component testing of gene-level heritability

One approach to test for gene-level heritability relies on a Wald test with the null hypothesis 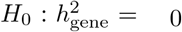, based on the point estimate and SE as derived in the above sections. To further improve power, we implemented an efficient variance component test in FLEX-h2 to test whether 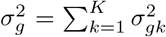 equals zero, which is equivalent to testing whether 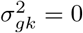 for all *K* annotations in the gene regions, given that 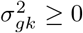. The genotypes in flanking regions are now treated as fixed effects in the covariates. We denote the conditional variance component tests as **FLEX-cond-test**.

Suppose the eigen-decomposition of ***K***_*i*_ is given by 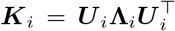, where **Λ**_*i*_ = diag(*λ*_*i*,1_,…, *λ*_*i,N*_) contains eigenvalues in decreasing order, and ***U*** _*i*_ is a unitary matrix. The test statistic is:

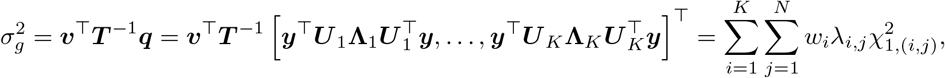

where ***v*** = [1,…, 1, 0]^⊤^, *w*_*i*_ = [***v*** ^⊤^ ***T*** ^−1^]_*i*_ represents the *i*-th entry, and 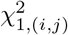 are i.i.d. chi-squared random variables with 1 degree of freedom. We perform a combined chi-square test using 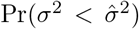, where 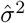 is the estimated 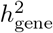 obtained via the Method-of-Moments approach. This test asks whether the observed 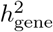 is larger than would be expected under the null hypothesis of no heritability contribution from that gene.

To approximate 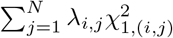, we employ the Satterthwaite approximation [45–47], which decomposes the weighted sum of chi-squared variables into two components 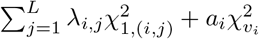. Here, *λ*_*i,j*_ denotes the eigenvalues of the variance component kernel associated with the *i*-th test, and 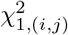 are independent chi-squared variables with 1 degree of freedom. The parameter *L* controls how many of the largest eigenvalue-weighted terms are retained explicitly, while the remaining contribution is captured by a single scaled chi-squared distribution with parameters *a*_*i*_ and *v*_*i*_, matched to preserve the first two moments of the omitted tail, with

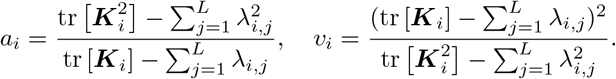

We empirically set *L* = 200, which we found to provide a good trade-off between computational efficiency and approximation accuracy across all tested settings. Additionally, we leverage the property that ***X***^⊤^ ***X*** and ***XX***^⊤^ share the same non-zero eigenvalues to accelerate computation. This is particularly beneficial because the number of variants *M* is much smaller than the number of individuals *N* in genotype data for a given gene region.

### Estimating gene-level heritability using summary statistics

Individual-level genotype data often remain inaccessible due to storage limitations and privacy concerns. To address this, we developed an extension of FLEX-h2 that operates on GWAS and burden test summary statistics, achieving accuracy equivalent to FLEX-h2 based on individual-level data. Following Jeong et al. [48], the left-hand side (***T***) of the normal equation (Equation 1) can be approximated using stratified stochastic LD scores (sketch-based approximations to LD scores) across annotations, while the right-hand side (***q***) is derived from GWAS and burden statistics. Asymptotic standard errors are obtained from an estimate of Cov[***q***], which we approximate using randomized trace estimation for computational efficiency as

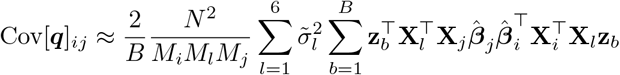

(see Supplemental Notes, “Derivation of the summary-statistics version of FLEX”, for derivations). We refer to this summary-statistics implementation as **FLEX-summ-h2** in the main analysis.

### Efficient estimation across multiple phenotypes

Notice that the matrix ***T***, which captures the trace terms involving genetic relatedness matrices, depends only on the genotypes and is independent of the phenotype vector ***y***. This structure enables computational reuse: once ***T*** is computed, it can be shared across traits for estimating gene-level heritability 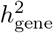. Let ***q***_*m*_ denote the vector of quadratic forms computed from the *m*th phenotype vector ***y***_*m*_, and let 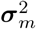 denote the corresponding vector of variance component estimates. Define ***Q*** = [***q***_1_ ***q***_2_ ··· ***q***_*M*_] and 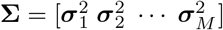 as matrices formed by stacking the per-phenotype vectors column-wise. Then, estimation of variance components across *M* phenotypes reduces to solving a single system:

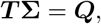

where each column of **Σ** contains the component estimates for a single trait. This formulation allows simultaneous estimation across multiple phenotypes using shared computational resources, yielding substantial efficiency gains, particularly in large-scale biobank settings.

### Genotype and phenotype data

We analyzed WES data from unrelated individuals of European ancestry in the UK Biobank. Sample-level quality control excluded individuals with excess genotype missingness, sex discrepancies between self-report and genetically inferred sex, or excess heterozygosity (defined as more than five standard deviations above the mean). Related individuals were identified and pruned to retain a maximal set of unrelated samples, resulting in a final sample of 348, 290 individuals. We then analyzed a subset of 153, 351 individuals from the second UK Biobank WES release. To improve genome coverage and facilitate conditional analyses on local LD, we merged imputed genotypes with WES variants, retaining WES variants over imputed variants when both were available. Imputed SNPs were filtered to exclude those with missingness *>* 1%, Hardy-Weinberg equilibrium *p <* 10^−7^, or INFO scores *<* 0.3. The final dataset comprised a total of 21, 512, 331 variants, including 11, 634, 606 imputed and 9, 877, 725 WES variants.

We applied FLEX to 20 quantitative traits, selected to span diverse biological systems, including anthropometry, blood pressure, lipid and glucose metabolism, liver, kidney, and pulmonary function. For phenotypes affected by medication, we applied trait-specific corrections (e.g., LDL direct cholesterol [49], systolic blood pressure [50]) prior to downstream analysis. To account for population structure and allele frequency-dependent stratification [25], we computed 10 principal components (PCs) within each of five MAF bins using ProPCA [51]: [0.01, 0.5), [0.01, 0.1), [10^−3^, 0.01), [10^−4^, 10^−3^), and [10^−5^, 10^−4^). Variants in the rarest bin (0, 10^−5^) were retained for analysis but excluded from PC computation due to high sampling noise. The final covariate model included sex, age, and 50 MAF-stratified PCs, totaling 52 covariates. Additional medication-related covariates were included for blood pressure traits (e.g., blood pressure medications, lipid-lowering agents, insulin, hormone replacement therapy, and oral contraceptives).

### Gene annotation

We used GENCODE release 46 to define gene boundaries [52], extending 10 kb upstream and downstream of each transcription start site (TSS) to capture proximal regulatory elements such as promoters and enhancers while minimizing inclusion of unrelated regions [53, 5]. Analyses were restricted to protein-coding genes. We excluded NOVEL genes due to limited external support and removed KNOWN genes lacking valid HGNC symbols to avoid ambiguity and ensure reproducibility. Genes with fewer than 100 variants after ultra-rare variant collapsing were also excluded to ensure stable estimation. After filtering, 18, 624 protein-coding genes remained for analysis.

### Variant annotation

Variants were annotated using the Ensembl Variant Effect Predictor (VEP v109). Loss-of-function (LoF) variants were defined according to high-confidence predictions from the LOFTEE plugin [3]. Missense variants were further characterized using established VEP-integrated predictive scores, including PolyPhen2-HumVar/HumDiv [54], SIFT [55], MutationTaster [56], and LRT [57]. To enable LD-aware analyses, we computed LD scores for each variant with respect to HapMap3 SNPs using GCTA [58]. LD scores were not computed against all genotyped variants to avoid bias due to uneven variant spacing. LD scores were used to partition variants within each MAF bin into low- and high-LD strata by median split for downstream stratified analyses.

### Gene set enrichment

To assess the biological relevance of trait-associated genes, we performed gene set enrichment analysis using pathway and regulatory annotations. For each trait, we identified a set of significant genes by applying a Bonferroni-corrected threshold of *p<* 0.05*/*18, 624, where 18, 624 denotes the total number of protein-coding genes tested. These trait-specific gene sets were then queried against the `g:Profiler` tool, which integrates multiple annotation sources, including GO biological processes, KEGG signaling and metabolic pathways, and transcription factor binding site data from TRANSFAC [59]. Enrichment significance was assessed using the built-in hypergeometric testing framework in `g:Profiler`, and multiple testing was controlled using the Benjamini-Hochberg false discovery rate (FDR) method. We report FDR-adjusted *p*-values for all significantly enriched terms.

### Overlap analysis

We obtained genome-wide association study (GWAS) summary statistics from the GWAS Catalog [12]. Trait names from our analysis were manually mapped to the most relevant traits in the catalog. A GWAS signal was considered significant if *p <* 5 *×* 10^−8^. We defined an overlap as occurring when a significant GWAS variant was mapped to the same gene as a gene-trait pair identified by FLEX (*p <* 0.05*/*18, 624). We recorded the number of unique GWAS hits per gene. Note that associated variants may be in linkage disequilibrium, and thus not statistically independent.

We also downloaded gene-based association statistics from the Genebass project [13], using mapped UK Biobank field IDs for trait harmonization. We considered *p*-values from the SKAT-O test applied to each of protein loss-of-function (pLoF), missense, and synonymous variants with MAF *<* 0.01. Gene-trait associations were deemed significant at *p<* 2.5 *×* 10^−7^.

### Simulation details

#### Calibration

We simulated phenotypes under a null model with no gene-level heritability using merged WES and imputed genotypes from chromosome 21. In these simulations, we selected a total of 117 genes with at least 100 variants after ultra-rare variant collapsing. We varied the background polygenic heritability, *i*.*e*., heritability attributed to variants outside of these genes,at levels 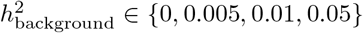. In the default setting, we set the proportion of causal variants to 10%. Following prior works [60, 61, 39], we specified the dependence between MAF and effect sizes by modeling the standardized genotype effect sizes such that their variance scaled with MAF *f* and LD score *w* as *w*^*a*^(*f* (1 − *f*))^*b*^ where *a* ∈ {0, 1} and *b* ∈ {0, 0.75}. To assess robustness to the causal SNP proportion, we fixed 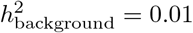 and varied the proportion of causal variants from 0.005 to 0.1. For each setting, we simulated 100 replicates.

#### Power and accuracy

To evaluate the ability of FLEX to detect gene-level heritability attributable to non-common variants, we designed simulations under a realistic genetic architecture where common variant effects are broadly polygenic and dispersed across the genome. Specifically, we fixed common variant heritability at 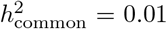 and introduced varying levels of gene-level heritability from non-common variants, with the sum ranging from 0.002 to 0.05. In each replicate, 5% of genes were randomly designated as causal, with non-common heritability apportioned across low-frequency, rare, and ultra-rare variants in a 4:4:2 ratio. To account for differing assumptions about how effect sizes scale with allele frequency, we drew SNP effects under both the uniform and MAF-dependent models.

#### Sensitivity to modeling parameters

To test the number of random vectors *B* in the trace approximation, we simulated phenotypes with *h*^2^ = 0.01 for both common and non-common components and applied FLEX using *B* = 10, 100, and 500. To assess the impact of LD flank size, we simulated phenotypes with 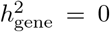 and varied intergenic background heritability over {0, 0.01, 0.1}. FLEX was applied with LD windows of 10 kb, 100 kb, and 1 Mb (100 replicates each).

#### Effect of ultra-rare variant collapsing

In the default simulations, ultra-rare variants were collapsed by aggregating effects within each functional category, with the variance for gene *g* and category 𝒱_*g*_ defined as 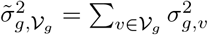. This assumes variants in the same category share a common effect size distribution. To test this, we also simulated phenotypes without collapsing, assigning 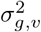 directly to each variant.

#### Runtime benchmarks

To evaluate scalability, we benchmarked FLEX across cohorts of 30, 000 to 100, 000 individuals and gene regions containing up to 2, 000 variants. Simulations were repeated 10 times per configuration, and compute time was recorded using wall-clock timing across 8 threads on a single core of an Intel Xeon Gold 6140 CPU (36 cores, 2.30 GHz base frequency).

#### Benchmarking calibration and power against existing methods

We benchmarked calibration and power for FLEX against two existing approaches, BHR and RARity. To assess calibration, we simulated phenotypes under a null model with no gene-level heritability, introducing background polygenic heritability in intergenic regions at levels 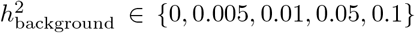. RARity was run with the authors’ recommended preprocessing: variants were filtered to MAC *>* 2 and pruned within 50 Mb sliding windows using an LD threshold of *r >* 0.9 [4]. BHR was excluded from comparisons at higher background heritability levels, as it is not designed to model common variants. To assess power and point estimation accuracy, we simulated phenotypes with total non-common 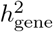 set to {0.01, 0.02, 0.05}. Phenotypes were simulated under the GCTA model within each MAF bin, and all methods were applied to estimate gene-level heritability stratified by MAF bin.

### Estimation of gene-level heritability in the UK Biobank

We estimated the gene-level heritability of each of the 18, 624 protein-coding genes across 20 quantitative traits in UKB. To account for local LD, estimation was conditioned on *±*100 kb regions flanking the defined gene boundaries. We used FLEX-h2 to estimate both point estimates and standard errors. To improve statistical precision for genes with marginal evidence of association (Wald test *p<* 0.05), we further applied a conditional variance component test (FLEX-cond-test).

### Estimation of exome-wide heritability in the UK Biobank

We applied FLEX-h2 to jointly estimate the phenotypic variance explained by coding variation across 18, 624 protein-coding genes (exome-wide heritability, 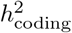) and that explained by genetic variants outside these genes. The model takes the form

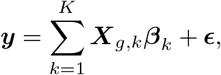

where ***X***_*g,k*_ is the standardized genotype matrix for the *k*-th annotation and *K* is the number of variant bins. We constructed a unified genotype matrix by concatenating variants from all genes (represented in the WES data) and intergenic regions (represented in the imputed SNPs), merging overlapping intervals across adjacent genes. Outside gene boundaries, we included only imputed variants with MAF *>* 10^−5^. As before, ***β***_*k*_ are vectors of *M*_*gk*_ entries which correspond to the SNP effect sizes in the *k*-th annotation while ***E*** is the *N* -vector of noise where 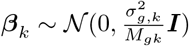, and 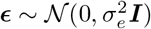.

To assess contributions from specific variant classes, we used two stratification schemes. The first partitioned variants into coding and intergenic regions, then into four MAF bins – common, low-frequency, rare, and ultra-rare – and further into high- and low-LD strata (*K* = 14; no ultra-rare variants in flanks). The second grouped variants into three MAF categories – common/low-frequency, rare, and ultra-rare – and stratified each by functional annotation (pLoF, missense, synonymous, other; *K* = 12). To estimate the heritability of a specific class of variants, *e*.*g*., rare coding variants, we sum the heritability estimates of the corresponding bins. We denote heritability from protein-coding variants by 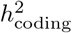 and that from all variants (both protein-coding and intergenic) by *h*^2^ and from sub-classes of variants analogously (so that 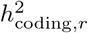 denotes heritability attributed to rare coding variants).

We defined the per-allelic effect size squared as

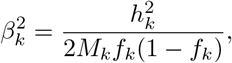

with standard error calculated as 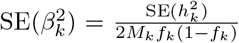, where 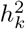represents the heritability explained by bin *k, M*_*k*_ is the number of variants within that bin, and *f*_*k*_ is the bin-specific mean MAF. To derive MAF-specific effect size estimates, we first summed heritability estimates across bins in that MAF range and then approximated their standard errors assuming they are independent.

To summarize per-trait estimates while accounting for variability and potential outliers, we used a robust random-effects meta-analysis framework [62, 63]. We first estimated the between-trait variance using the DerSimonian-Laird method [64] and then applied an iterative procedure with Huber weighting [65] to reduce the influence of traits with unusually large or inconsistent effect sizes. This approach balances statistical efficiency with robustness to outliers.

## Acknowledgments

This research was conducted using the UK Biobank Resource under application 331277. We thank the participants of UK Biobank for making this work possible. This work was funded, in part, by NIH grants R35GM153406 (Z.L., B.F., M.J., P.A., A.A., and S.S.), R01HL170604 (P.P.), R01DK132775 (P.P.), and NSF grant CAREER-1943497 (S.S.).

## Data Availability

The UK Biobank dataset is available upon application at https://www.ukbiobank.ac.uk/. The Genebass database is freely accessible at https://app.genebass.org/. The g:Profiler tool can be accessed at https://biit.cs.ut.ee/gprofiler/. The GWAS Catalog is available at https://www.ebi.ac.uk/gwas/. Gene annotations can be downloaded from https://www.gencodegenes.org/human/release_46lift37.html.

## Code Availability

All software and code used in this study are publicly available: GCTA (https://yanglab.westlake.edu.cn/software/gcta), PLINK (https://www.cog-genomics.org/plink/2.0), BHR (https://github.com/ajaynadig/bhr), RARity (https://github.com/GMELab/RARity), ProPCA (https://github.com/sriramlab/ProPCA), and FLEX (https://github.com/sriramlab/GENIE/tree/main/FLEX).

