## Supplemental Information for "Comprehensive gene heritability estimation reveals the genetic architecture of rare coding variants underlying complex traits"

### Supplemental Notes

#### S1 Derivation of the summary-statistics version of FLEX

As shown in Jeong et al. [48], the left-hand side ($\boldsymbol{T}$) of Equation 1 can be expressed in terms of stratified LD scores, while the right-hand side ($\boldsymbol{q}$) can be computed from GWAS and burden test statistics. For the asymptotic variance, we require $\mathrm{Cov}[\boldsymbol{q}]$, where the $(i, j)$ entry is

$$\begin{aligned}\mathrm{Cov}[\mathbf{q}]_{ij} &\approx 2\sum_{l=1}^{6} \tilde{\sigma}_l^2\, \mathbf{y}^\top \mathbf{K}_i \mathbf{K}_l \mathbf{K}_j \mathbf{y} \\ &= 2\sum_{l=1}^{6} \tilde{\sigma}_l^2\, \frac{N^2}{M_i M_l M_j}\, \hat{\boldsymbol{\beta}}_i^\top \mathbf{X}_i^\top \mathbf{X}_l \mathbf{X}_l^\top \mathbf{X}_j \hat{\boldsymbol{\beta}}_j,\end{aligned}$$

where $\hat{\boldsymbol{\beta}}_i = \mathbf{X}_i^\top \mathbf{y}/N$ denotes the GWAS effect size vector for annotation $i$, $\mathbf{X}_i$ is the standardized genotype matrix, and $M_i$ is the number of variants in annotation $i$.

To approximate the quadratic form efficiently, we use the Hutchinson trace estimator, $\mathbb{E}[\mathrm{tr}(\mathbf{A})] = \mathbb{E}[\mathbf{z}^\top \mathbf{A}\mathbf{z}]$ for $\mathbf{z} \sim \mathcal{N}(0, \mathbf{I}_N)$. Applying this identity yields:

$$\begin{aligned}\mathbb{E}_{\mathbf{z}}[\hat{\boldsymbol{\beta}}_i^\top \mathbf{X}_i^\top \mathbf{X}_l \mathbf{X}_l^\top \mathbf{X}_j \hat{\boldsymbol{\beta}}_j] &= \mathbb{E}[\mathrm{tr}(\hat{\boldsymbol{\beta}}_i^\top \mathbf{X}_i^\top \mathbf{X}_l \mathbf{X}_l^\top \mathbf{X}_j \hat{\boldsymbol{\beta}}_j)] \\ &= \mathbb{E}[\mathrm{tr}(\mathbf{X}_l^\top \mathbf{X}_j \hat{\boldsymbol{\beta}}_j \hat{\boldsymbol{\beta}}_i^\top \mathbf{X}_i^\top \mathbf{X}_l)] \\ &= \mathbb{E}_{\mathbf{z}}[\mathbf{z}^\top \mathbf{X}_l^\top \mathbf{X}_j \hat{\boldsymbol{\beta}}_j \hat{\boldsymbol{\beta}}_i^\top \mathbf{X}_i^\top \mathbf{X}_l \mathbf{z}] \\ &\approx \frac{1}{B}\sum_{b=1}^{B} \mathbf{z}_b^\top \mathbf{X}_l^\top \mathbf{X}_j \hat{\boldsymbol{\beta}}_j \hat{\boldsymbol{\beta}}_i^\top \mathbf{X}_i^\top \mathbf{X}_l \mathbf{z}_b,\end{aligned}$$

where $\{\mathbf{z}_b\}_{b=1}^B$ are standard normal random vectors. By precomputing the sketched matrices $\mathbf{X}_i^\top \mathbf{X}_l \mathbf{z}_b$, we greatly reduce computation time when estimating $\mathrm{Cov}[\boldsymbol{q}]$ across multiple annotations and traits.

#### S2 Impact of linkage disequilibrium on gene-level heritability estimation

While the simulations in the main text demonstrated the robustness of FLEX to LD-induced confounding when conditioning on flanking regions, here we seek to systematically quantify the factors that impact the magnitude of LD-related inflation when such conditioning is omitted. Linkage disequilibrium between gene-body variants and surrounding regions is a well-recognized source of bias in local heritability estimation [25, 5], particularly for short genes or those located in high-LD regions.

To directly assess the extent of LD-induced inflation, we performed simulations under the null model of no gene-level heritability, varying key parameters – average LD score and gene size – while simulating background polygenic heritability (excluding gene regions) at levels ranging from 0 to 0.1. Simulations were restricted to protein-coding genes on chromosome 21, selecting representative genes with maximum, median, and minimum values for each parameter of interest while controlling for the other (40th – 60th percentile). Gene-level heritability was then estimated using FLEX without LD conditioning, across 10 replicates for each scenario.

Clear inflation in $h^2_{\text{gene}}$ estimates was observed for shorter genes. For example, the small gene *ATP5PO* (426 SNPs) exhibited inflated estimates of $h^2_{\text{gene}} = 4.80 \pm 1.80 \times 10^{-4}$ when $h^2_{\text{background}} = 0.1$, while the larger gene *SLC37A1* (1,661 SNPs) remained near zero (Figure S24a). Similarly, genes with elevated LD scores exhibited greater inflation. For instance, *JAM2* (LD score = 8.33) produced $h^2_{\text{gene}} = 5.53 \pm 4.08 \times 10^{-4}$ and $10.9 \pm 6.65 \times 10^{-4}$ at $h^2_{\text{background}} = 0.05$ and 0.1, respectively. In contrast, genes with lower LD scores, such as *ZBTB21* (2.25) and *KRTAP10-11* (3.85), displayed no appreciable inflation (Figure S24b).

We further examined how LD confounding varies across allele frequency spectra. Using $h^2_{\text{background}} = 0.1$, gene-level heritability was estimated for all 117 genes on chromosome 21 without LD conditioning. Averaged over 10 replicates, inflation was observed for rare, low-frequency, and common variants in genes with high LD scores ($> 10$), whereas ultra-rare variant estimates remained largely unaffected. Specifically, mean $h^2_{\text{gene}}$ estimates for ultra-rare, rare, low-frequency, and common variants were $8.32 \times 10^{-7}$ (s.e. $1.18 \times 10^{-5}$), $5.99 \pm 5.00 \times 10^{-5}$, $2.48 \pm 1.55 \times 10^{-4}$, and $2.47 \pm 1.20 \times 10^{-4}$, respectively (Figure S25a). Similarly, when stratifying by gene size, short genes (fewer than 200 variants) showed upward bias for all MAF bins except ultra-rare variants (Figure S25b). These results demonstrate that LD tagging of neighboring non-causal variants can artifactually inflate gene-level heritability estimates, particularly for smaller genes and those located in regions of high LD.

#### Supplemental Figures

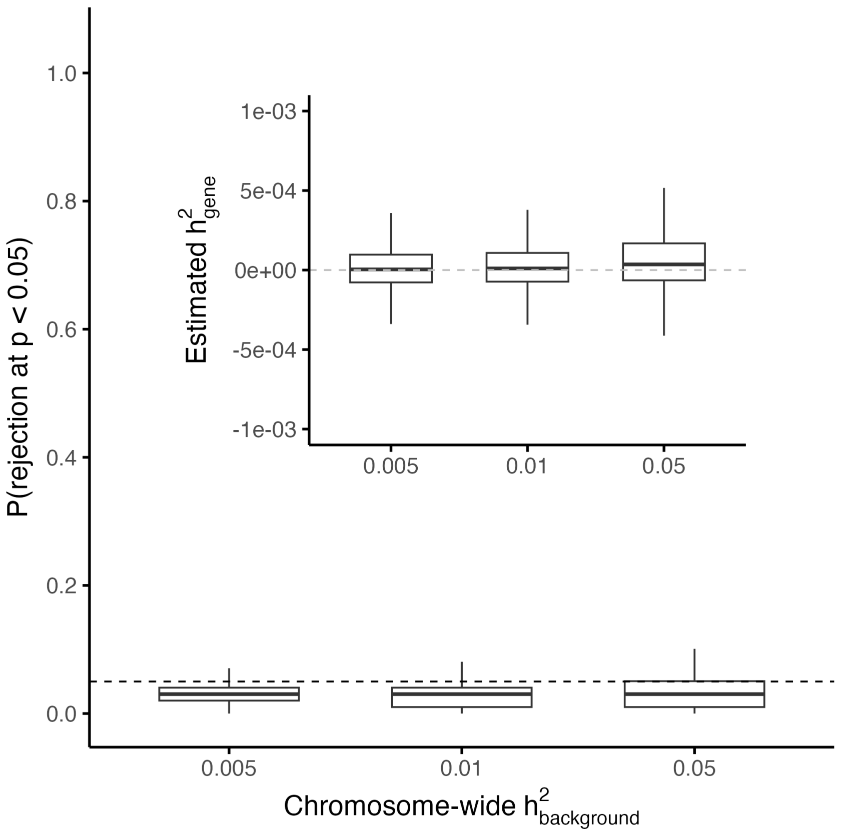


Figure S1: **Calibration of FLEX under LDAK simulations.** False positive rate, defined as the proportion of tests with $p < 0.05$ under null simulations ($h^2_{\text{gene}} = 0$), with the inset showing the distribution of estimated $h^2_{\text{gene}}$. Chromosome-wide $h^2_{\text{background}}$ was varied in LDAK simulations, where the variance of standardized genotype effect sizes was proportional to the LD score $w$ and MAF $f$ as $w^a(f(1-f))^b$, with $a = 1$ and $b = 0.75$, within each MAF bin.

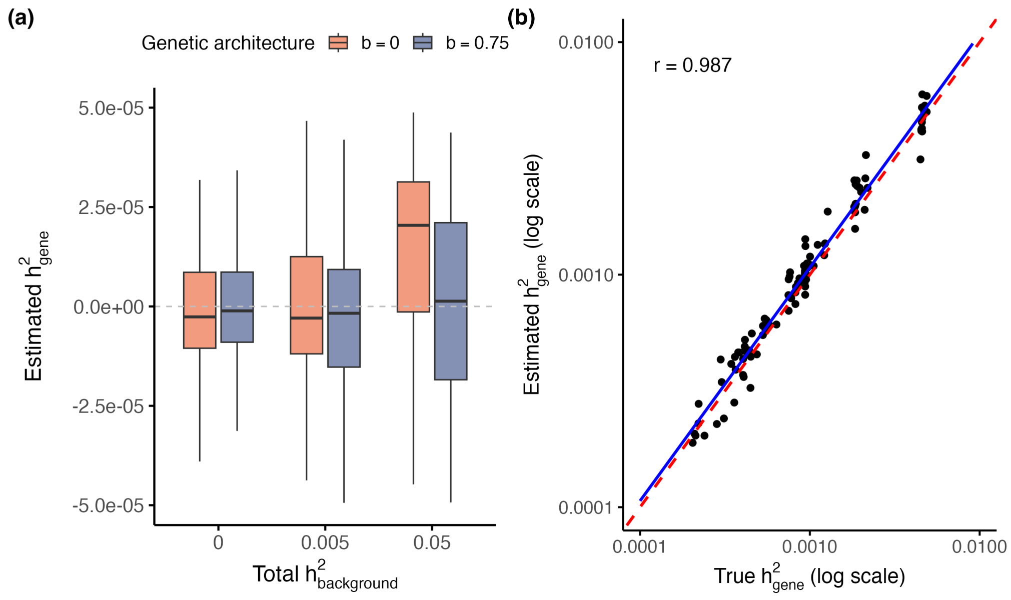


Figure S2: **Calibration and statistical power of the summary-based version of FLEX.** **(a)** Distribution of estimated $h^2_{\text{gene}}$ for genes with no causal variants, under varying $h^2_{\text{background}}$ and genetic architectures (the variance of standardized genotype effect sizes proportional to $(\text{MAF}(1-\text{MAF}))^b$ where $b \in \{0, 0.75\}$). FLEX-summ-h2 remains unbiased under both architectures. **(b)** Comparison of estimated versus true $h^2_{\text{gene}}$ on the log scale across simulations with nonzero gene-level heritability. FLEX-summ-h2 exhibits strong concordance with the truth (Pearson $r = 0.987$).

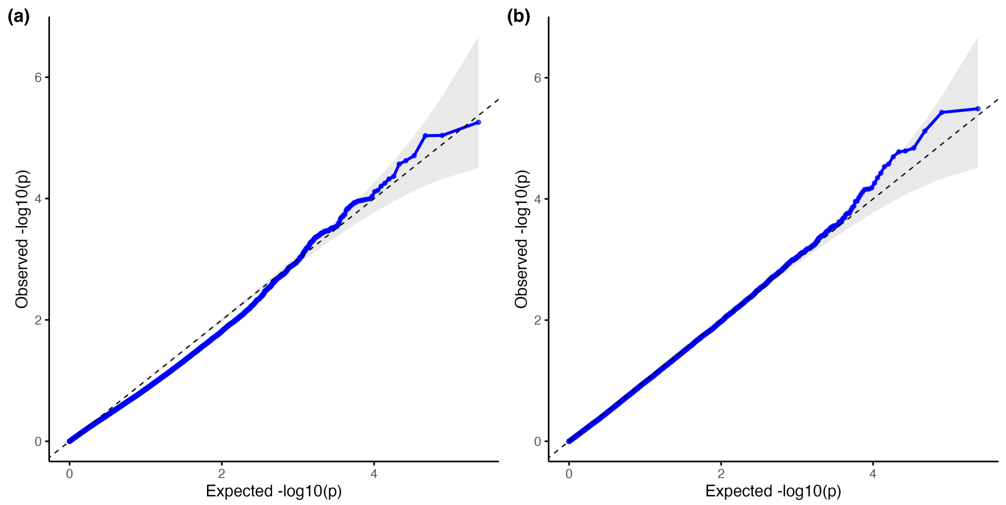


Figure S3: **QQ plots showing calibration of tests in FLEX.** Pooled $p$-values across null simulation scenarios are shown for **(a)** the Wald test in FLEX-h2 and **(b)** FLEX-cond-test. Both tests exhibit good calibration under the null, with the Wald test showing slight conservativeness at small $p$-values.

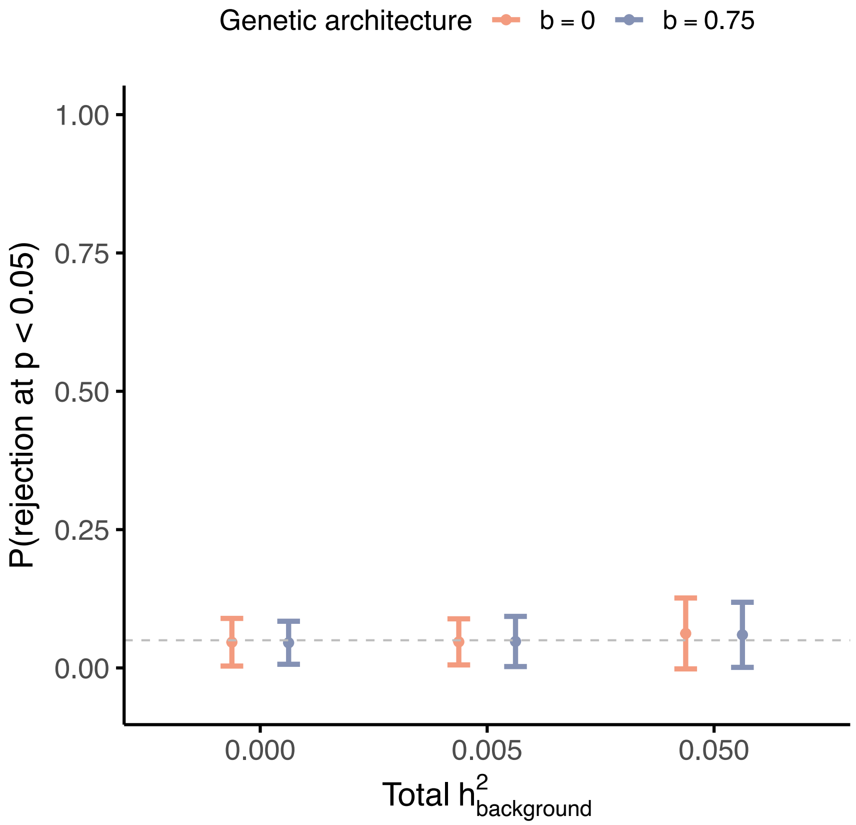


Figure S4: **Calibration of the conditional variance component test.** FPR defined as the proportion of $p < 0.05$ under null simulations ($h^2_{\text{gene}} = 0$), shown as mean $\pm$ 95% confidence intervals across genes. Results are shown for $h^2_{\text{background}} \in \{0, 0.005, 0.05\}$ and simulations under different genetic architectures.

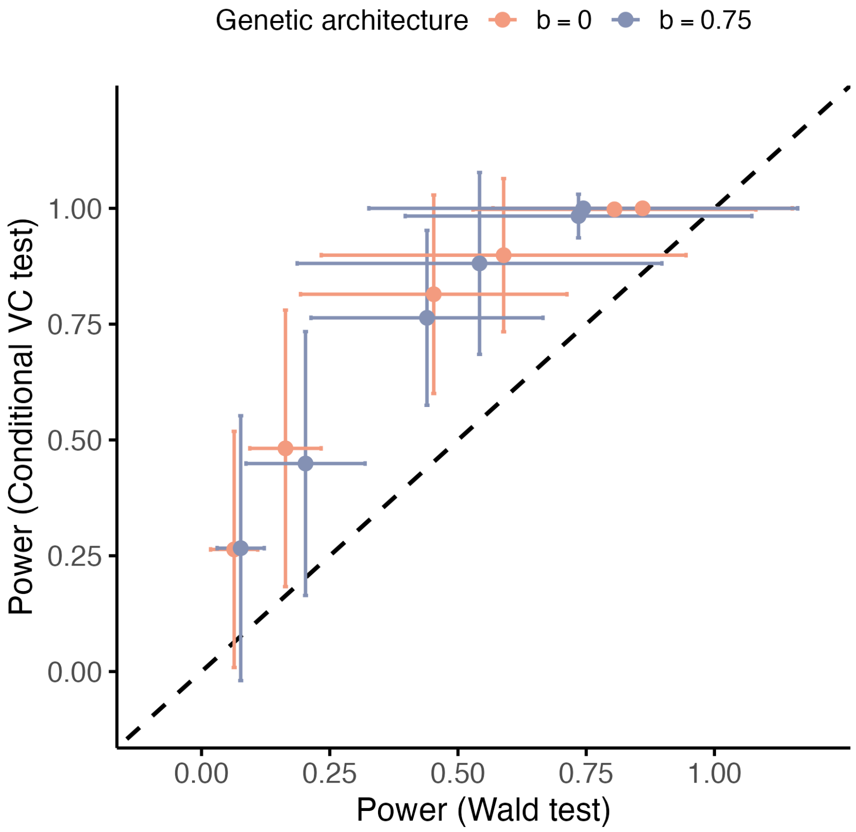


Figure S5: **Power comparison between conditional variance component and the Wald tests.** Power to detect gene-level heritability across a range of true simulated values using either the conditional variance component test or the Wald test. The conditional test consistently provides higher power, particularly for moderate heritability levels, under both genetic architectures.

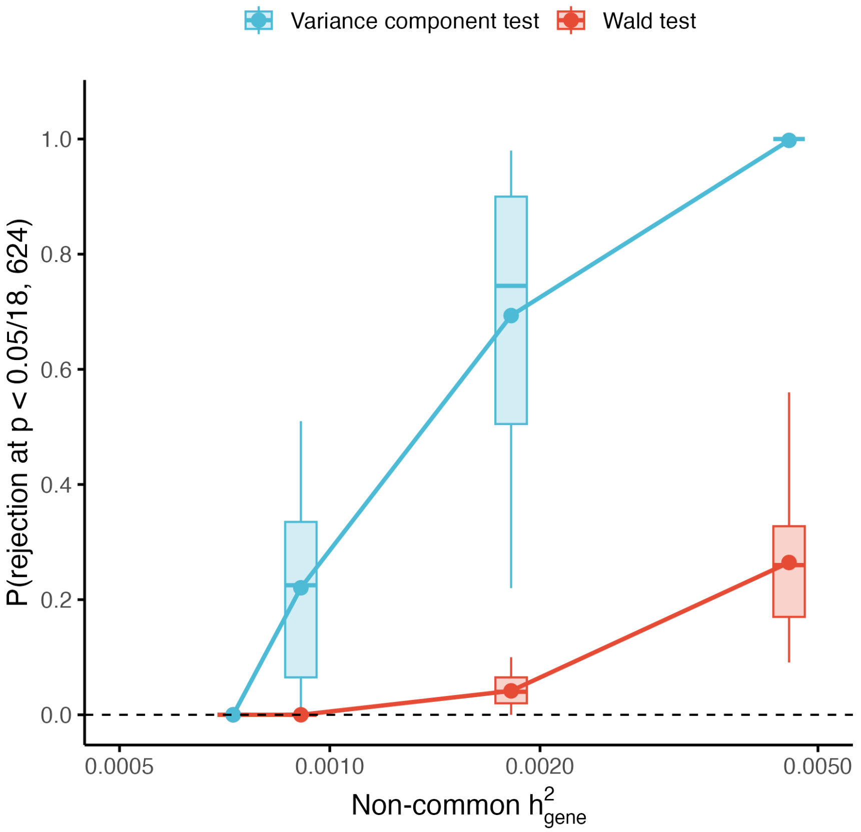


Figure S6: **Genome-wide power comparison between conditional variance component and the Wald tests.** Power to detect $h^2_{\text{gene}}$ as a function of non-common $h^2_{\text{gene}}$ at the genom-wide threshold of $p < 0.05/18,624$. The $x$-axis denotes the simulated non-common $h^2_{\text{gene}}$ for causal genes. Boxplots show results across 100 replicates per setting under different genetic architectures, with the dashed line indicating the Bonferroni-corrected threshold of $0.05/18,624$.

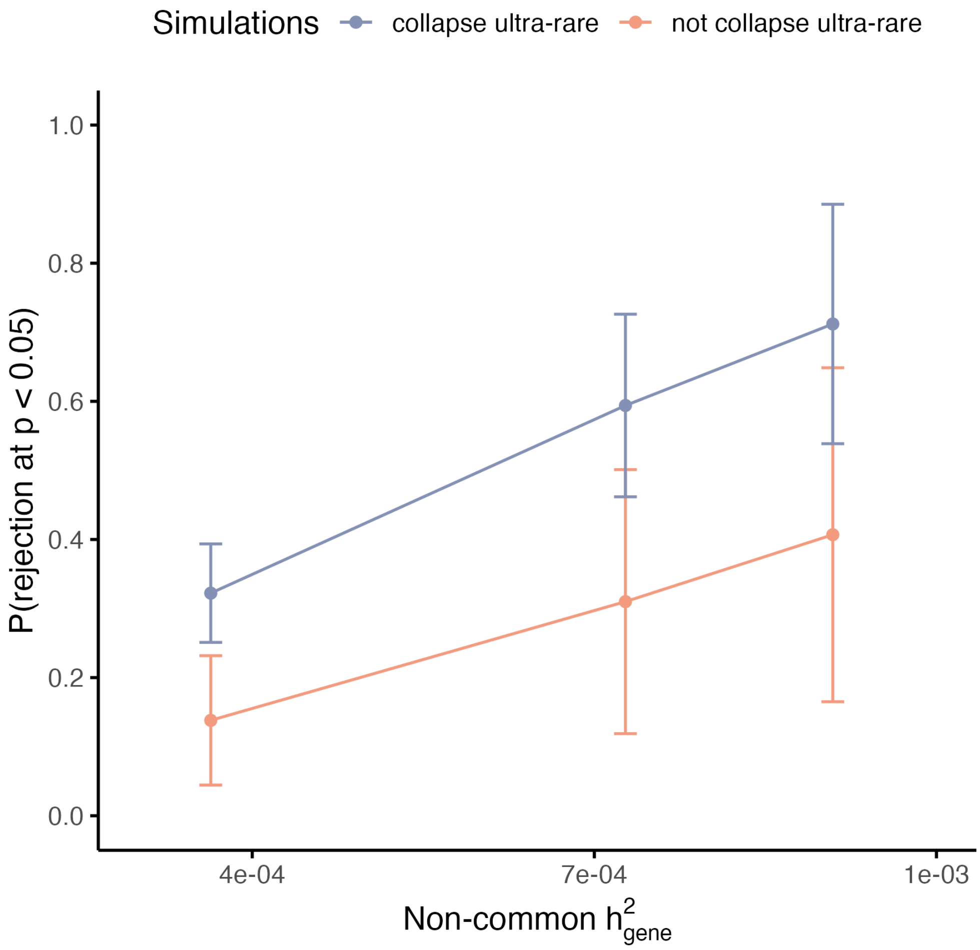


Figure S7: **Misspecification of simulations without collapsing ultra-rare variants.** Selected simulation settings were repeated without collapsing ultra-rare variants. In these scenarios, FLEX exhibited reduced power.

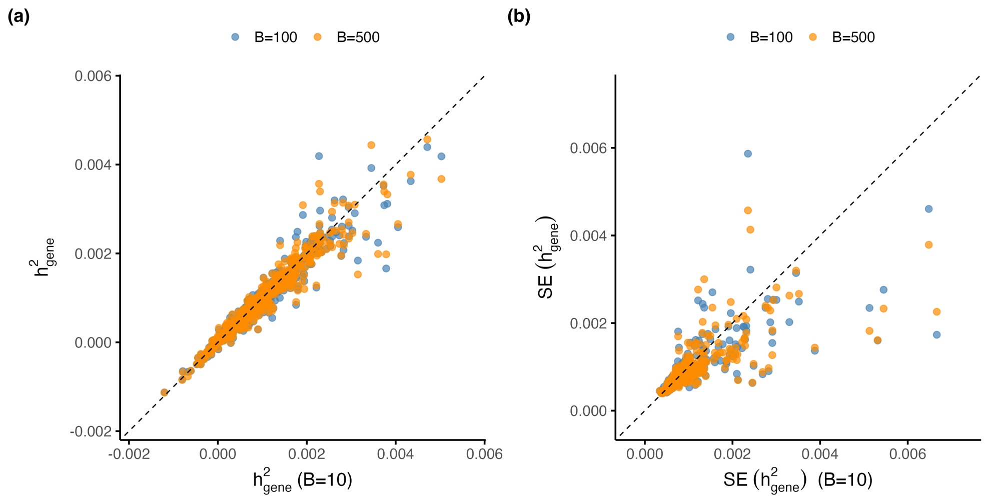


Figure S8: **Concordance of heritability estimates across sketch sizes.** Comparison of FLEX-estimated **(a)** $h^2_{\text{gene}}$ and **(b)** standard errors using different numbers of random vectors ($B = 10$, 100, 500) for trace estimation.

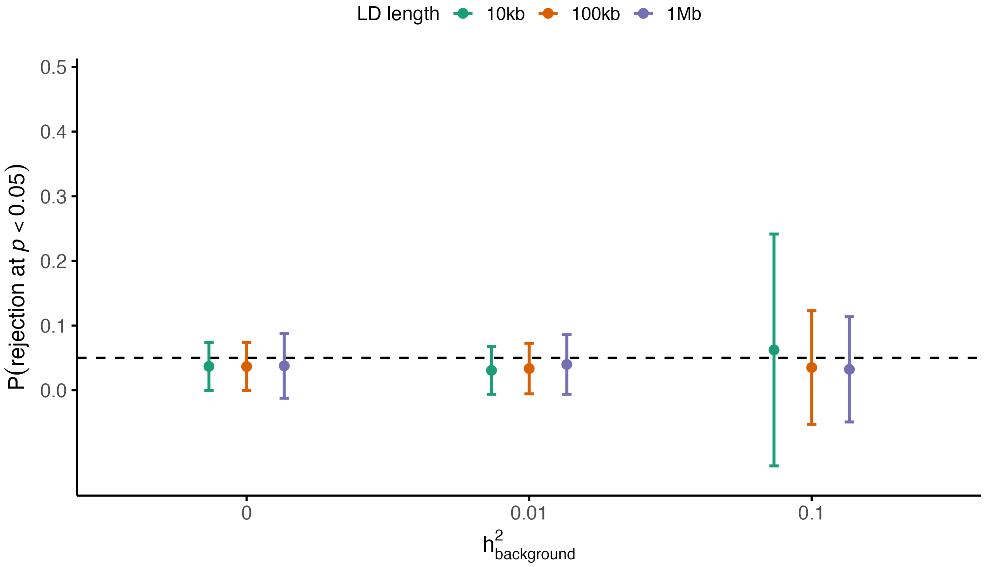


Figure S9: **Effect of flanking LD window size on false positive rate in calibration.** FPR defined as the proportion of $p < 0.05$ under null simulations ($h^2_{\text{gene}} = 0$), shown as mean $\pm$ 95% confidence intervals across genes. Results are stratified by background heritability ($h^2_{\text{background}}$) and LD conditioning window size (10 kb, 100 kb, 1 Mb). All window sizes yield FPR close to the nominal 0.05 level. However, at $h^2_{\text{background}} = 0.1$, the 10 kb window shows greater uncertainty in FPR compared to 100 kb and 1 Mb, supporting the use of 100 kb as the default LD correction window.

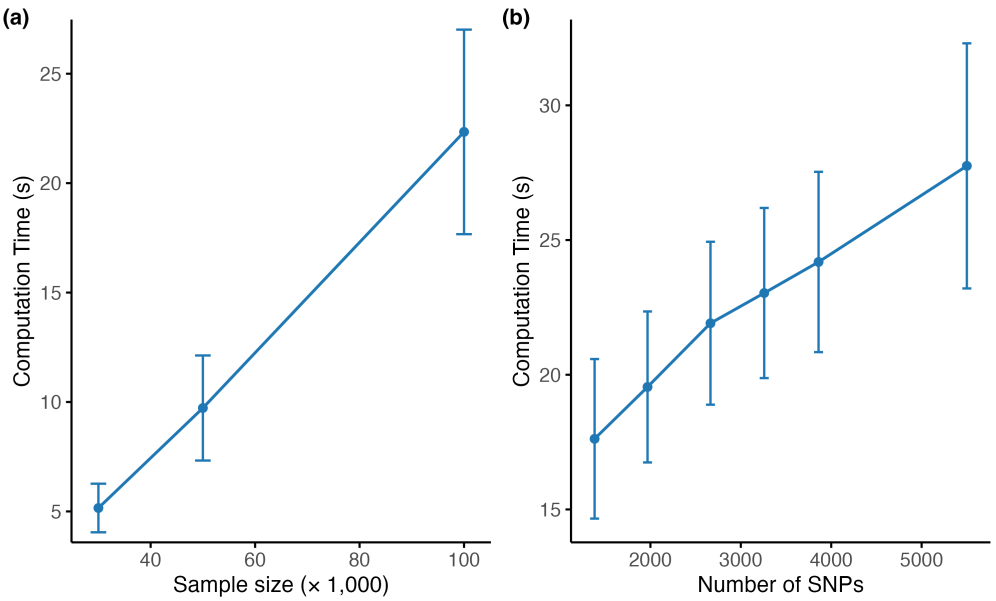


Figure S10: **Runtime scaling of FLEX.** **(a)** Mean runtime (± standard deviation) for FLEX as a function of sample size, evaluated across 117 genes on chromosome 21. **(b)** Runtime increases approximately linearly with the number of SNPs per gene. We performed the benchmark on genotypes of 100K individuals. Genes on chromosome 21 were grouped into deciles based on LD-pruned SNP count using quantile binning. All benchmarks were conducted using a single core of an Intel Xeon Gold 6140 CPU (36 cores, 2.30 GHz base frequency).

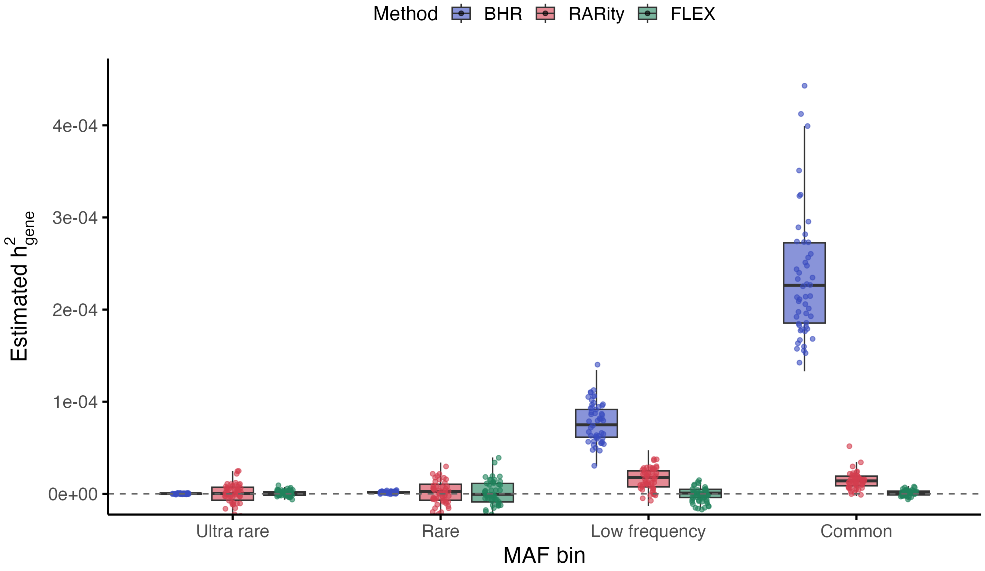


Figure S11: **Comparison of gene-level heritability estimates across MAF bins under the null.** Estimated $\hat{h}^2_{\text{gene}}$ across MAF bins for 117 genes with no causal variants, under a baseline background heritability of $h^2_{\text{background}} = 0.01$. FLEX remains well-calibrated across all MAF bins, with median estimates tightly centered near zero for common and low-frequency variants. RARity shows mild inflation in these bins, while BHR exhibits substantial upward bias, particularly for common variants. All methods yield approximately null-centered estimates for rare and ultra-rare variants.

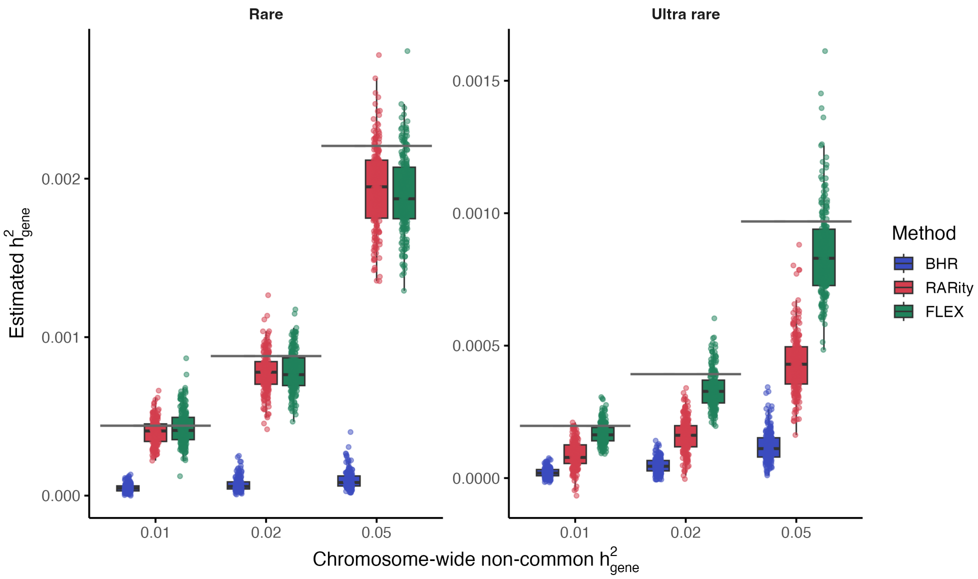


Figure S12: **Comparison of methods for estimating gene-level heritability under rare variant architectures.** Estimated $h^2_{\text{gene}}$ across increasing levels of simulated non-common variant heritability ($h^2_{\text{gene}} \in \{0.01, 0.02, 0.05\}$), evaluated separately for rare (left) and ultra-rare (right) variants. All methods were well-calibrated under the null and compared here under nonzero signal with 5% of genes designated as causal and background heritability fixed at $h^2_{\text{background}} = 0.01$. FLEX (green) consistently recovered a greater fraction of the simulated heritability and exhibited lower variance, while RARity (red) underestimated ultra-rare signal due to exclusion of singletons and LD pruning. BHR (blue) captured only a small fraction of the heritability across both bins.

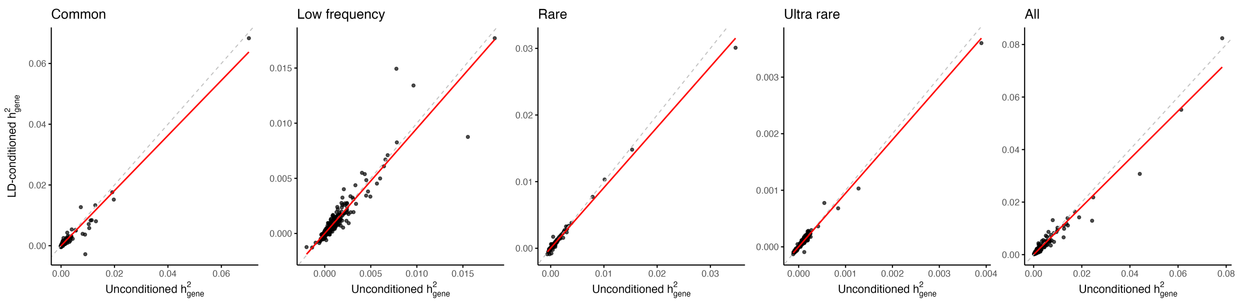


Figure S13: **Comparison of per-SNP heritability before and after LD conditioning across MAF bins.** Each point represents a gene–trait pair that is marginally significant ($p < 0.05$). The $x$- and $y$-axes show estimates of $h^2_{\text{gene}}$ without and with LD conditioning, respectively. The dotted diagonal denotes $y = x$, and the red line indicates the best-fit linear regression. Across all MAF bins except ultra-rare, LD conditioning significantly reduced per-SNP heritability estimates (one-sided $t$-test). Specifically, we observed significant downward shifts for common ($p = 5.84 \times 10^{-8}$), low-frequency ($p = 2.21 \times 10^{-5}$), and rare ($p = 2.45 \times 10^{-6}$) variants. Estimates for ultra-rare variants were largely unchanged ($p = 0.059$).

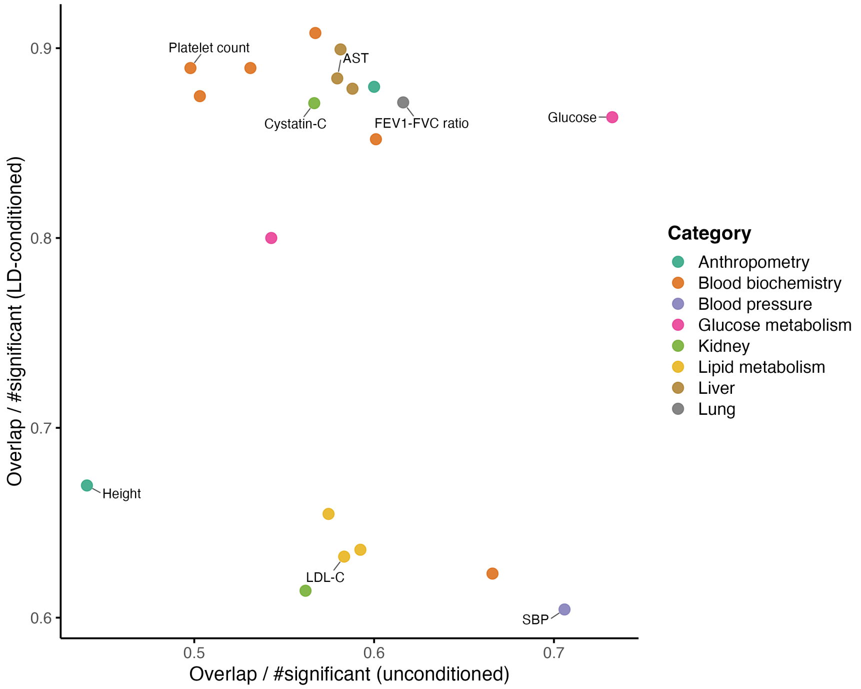


Figure S14: **Overlap of significant genes before and after LD conditioning.** Each point represents a trait. We identified significant genes under both unconditioned and LD-conditioned models using a threshold of $p < 0.05$. Overlap is defined as the number of genes that are significant in both models. The $x$- and $y$-axes show the fraction of overlapping genes relative to the unconditioned and conditioned sets, respectively.

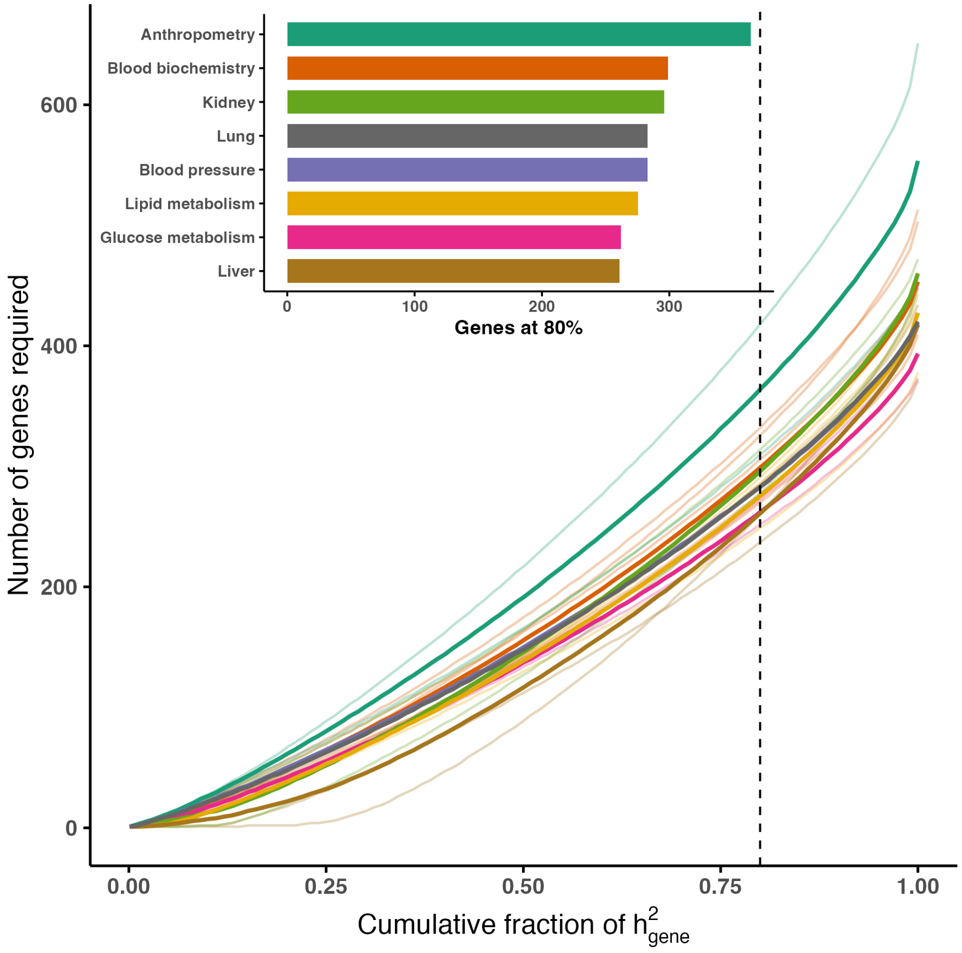


Figure S15: **Cumulative contribution of genes to rare and ultra-rare variant heritability.** Each curve shows the number of genes required to explain a given cumulative fraction of gene-level heritability ($h^2_{\text{gene}}$) based on rare and ultra-rare variants, for an individual trait (faint lines) or trait category (solid lines). The vertical dashed line indicates the point at which 80% of total $h^2_{\text{gene}}$ is captured. The inset summarizes the number of genes required to reach this 80% threshold across trait categories, providing a comparative view of polygenicity.

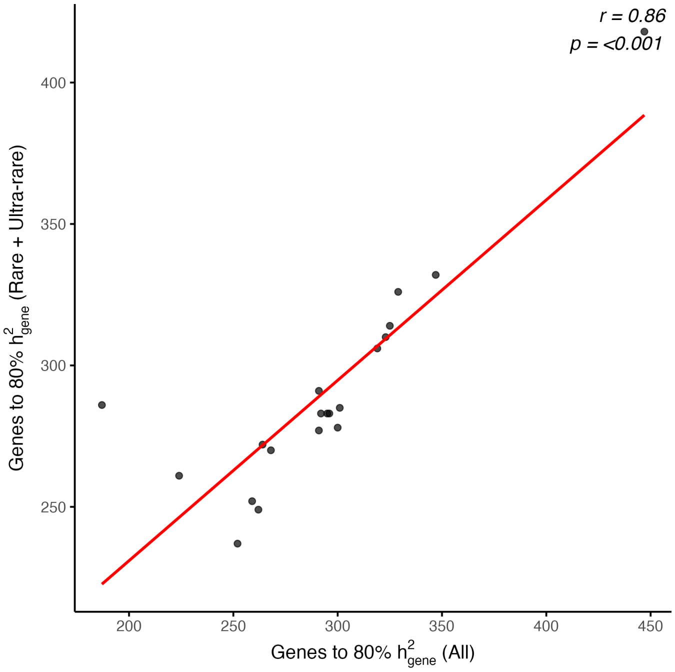


Figure S16: **Polygenicity of rare versus all variants.** Scatterplot shows the number of genes required to explain 80% of genome-wide $h^2_{\text{gene}}$ using all variants (x-axis) versus rare + ultra-rare variants (y-axis) across traits. Red line indicates linear regression fit. Annotated with Pearson $r = 0.858$ and $p = 1.30 \times 10^{-6}$.

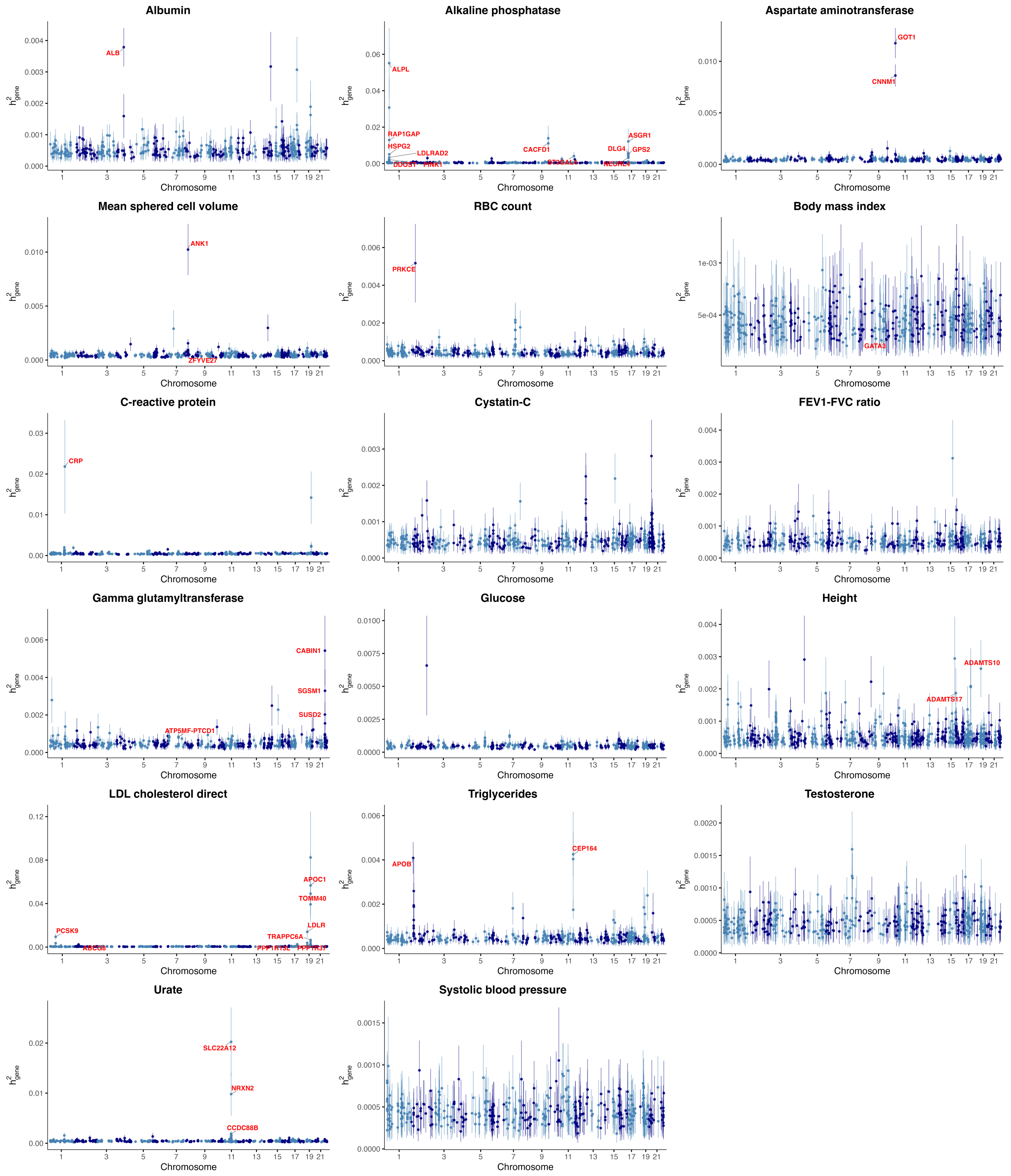


Figure S17: **Gene-level heritability of each trait.** Genome-wide Manhattan-style plots for gene-level heritability estimates of all 20 traits except platelet count, hemoglobin A1c, and HDL cholesterol. Each point represents a gene, with vertical lines indicating standard errors of the heritability estimates. Genes with genome-wide significant heritability ($p < 0.05/18,624$) are labeled in red. Note that some gene regions overlap due to the 10 kb extension of gene boundaries to include proximal regulatory regions (e.g., *APOC1* and *TOMM40*).

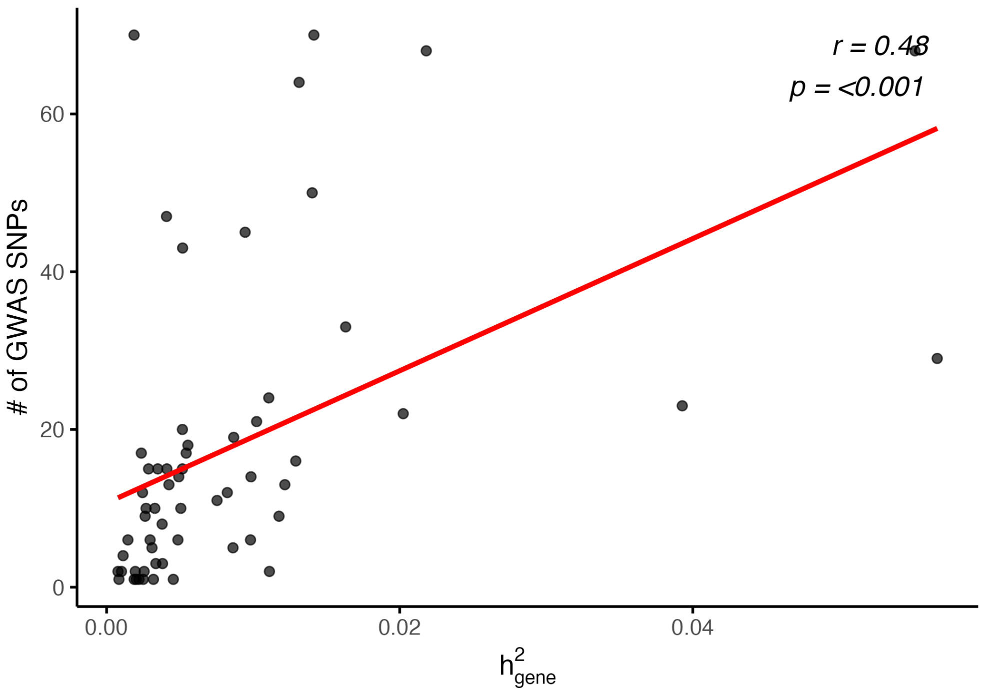


Figure S18: **Relationship between gene-level heritability significance and the number of GWAS loci.** We examined genome-wide significant gene–trait pairs from FLEX and quantified the number of GWAS loci (SNPs with $p < 5 \times 10^{-8}$) from the GWAS Catalog per gene–trait pair. The x-axis shows FLEX $p$-values, and the y-axis shows the corresponding GWAS SNP count. A positive correlation was observed ($r = 0.48$, $p = 3.21 \times 10^{-5}$), suggesting that FLEX significance is often supported by polygenic GWAS signal.

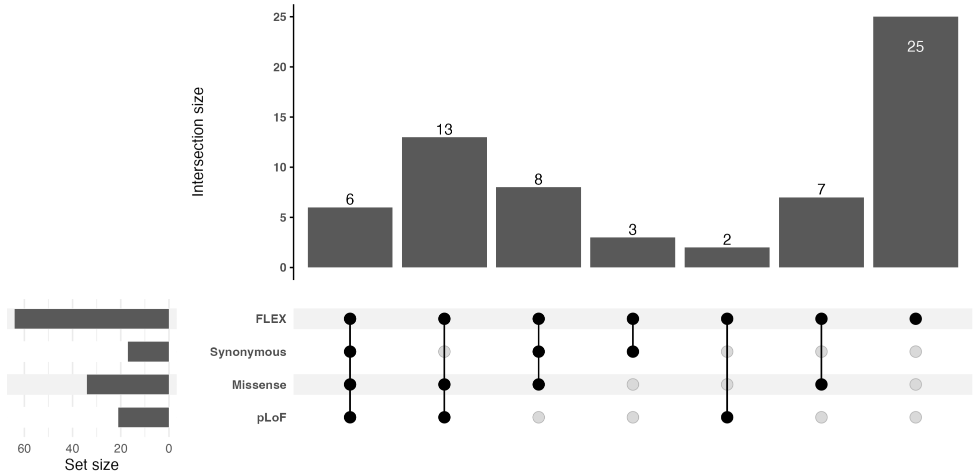


Figure S19: **Overlap of significant gene–trait pairs across functional bins.** UpSet plot visualizing the intersection of significant gene–trait pairs from FLEX and gene-level associations across variant classes (pLoF, missense, and synonymous). FLEX significance was defined at $p < 0.05/18, 624$, while Genebass gene-based associations used a threshold of $p < 2.5 \times 10^{-7}$. The top panel displays the number of overlapping gene–trait pairs for each combination of functional annotations, and the bottom panel shows the total number of significant pairs per method. Only pairs passing the respective significance thresholds for a given trait were included.

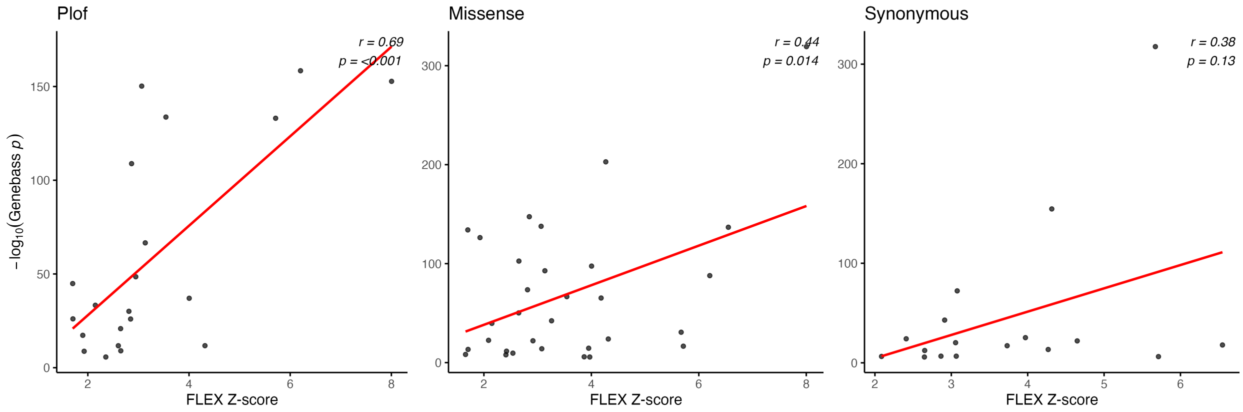


Figure S20: **Concordance between FLEX and Genebass burden test results.** Scatterplots compare gene-level $z$-scores from FLEX, defined as $h^2_{\text{gene}}/\text{SE}(h^2_{\text{gene}})$, against $-\log_{10}(p)$ values from Genebass SKAT-O tests across pLoF, missense, and synonymous variants. Each point represents a gene–trait pair. Using Genebass's recommended significance threshold ($p < 2.7 \times 10^{-7}$), we identified 21, 34, and 17 overlapping gene–trait pairs for pLoF, missense, and synonymous variants, respectively. Among these, SKAT-O $p$-values were significantly correlated with gene-level $z$-scores for pLoF ($r = 0.694$, $p = 4.79 \times 10^{-4}$) and missense ($r = 0.437$, $p = 0.014$) variants, but not for synonymous variants ($p = 0.130$).

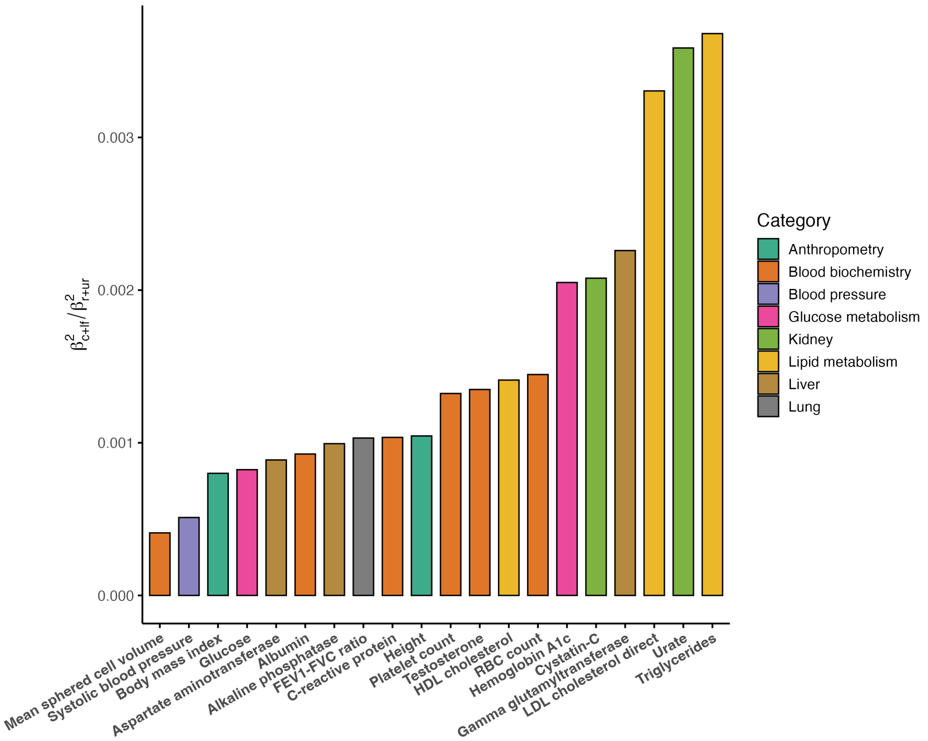


Figure S21: **Relative per-allelic effect sizes of rare versus common variants.** Each bar represents a trait, colored by category, showing the ratio of per-allelic effect sizes ($\alpha_{\text{maf}} = \beta^2_{c+lf}/\beta^2_{r+ur}$) for rare and ultra-rare ($\beta^2_{r+ur}$) versus common and low-frequency variants ($\beta^2_{c+lf}$). Traits with larger values exhibit stronger rare variant enrichment.

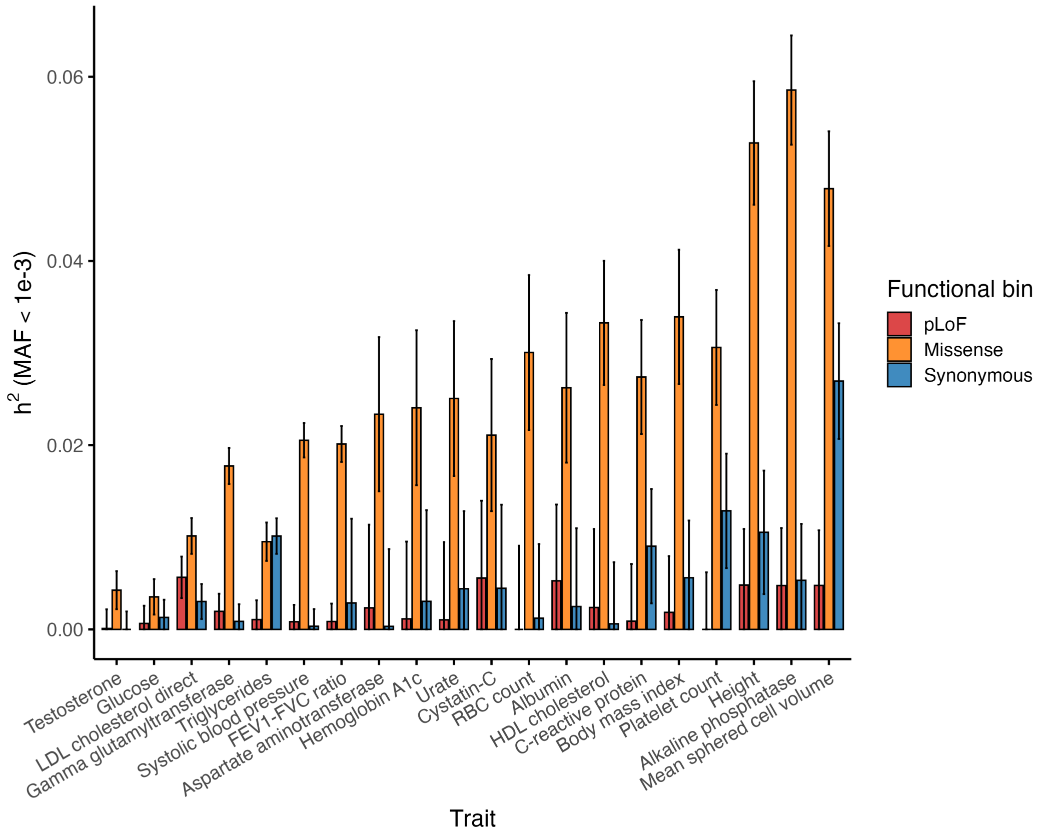


Figure S22: **Genome-wide heritability estimates across functional annotations.** Error bar plots of heritability estimates of pLoF, missense, and synonymous variants (MAF $\leq 10^{-3}$) across traits.

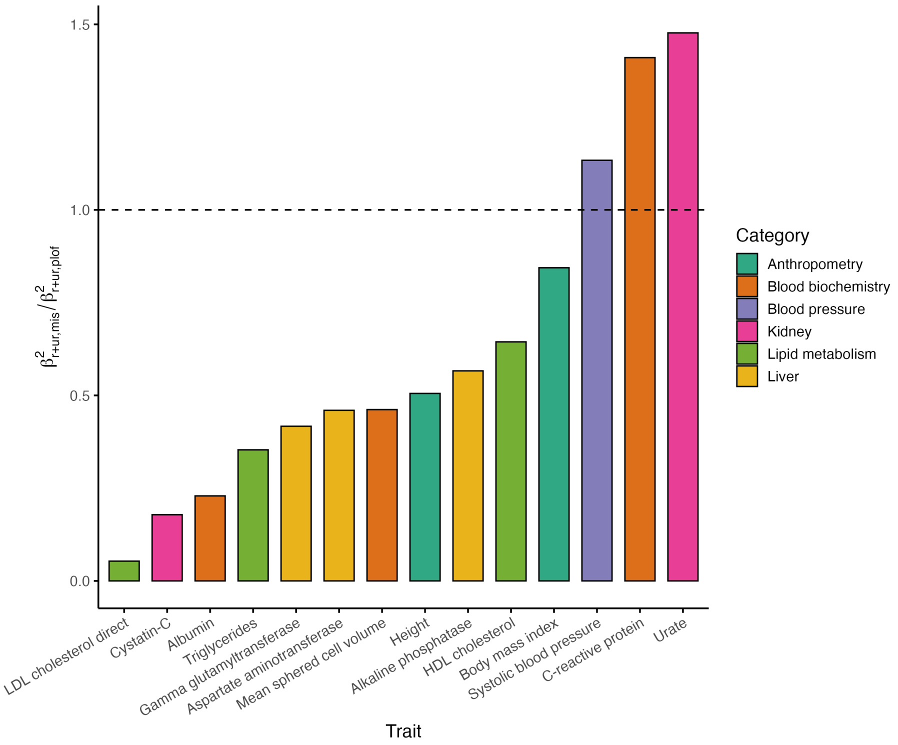


Figure S23: **Relative per-allele effect sizes of rare missense versus pLoF variants.** Each bar represents a trait, colored by category, and shows the ratio of per-allele squared effect sizes ($\alpha_{\text{func}} = \beta^2_{r+ur,\text{mis}}/\beta^2_{r+ur,\text{plof}}$) for rare and ultra-rare missense variants ($\beta^2_{r+ur,\text{mis}}$) relative to pLoF variants ($\beta^2_{r+ur,\text{plof}}$). Only traits with valid positive ratios are shown. Three traits exhibited stronger missense effects than pLoF effects ($\alpha_{\text{func}} > 1$).

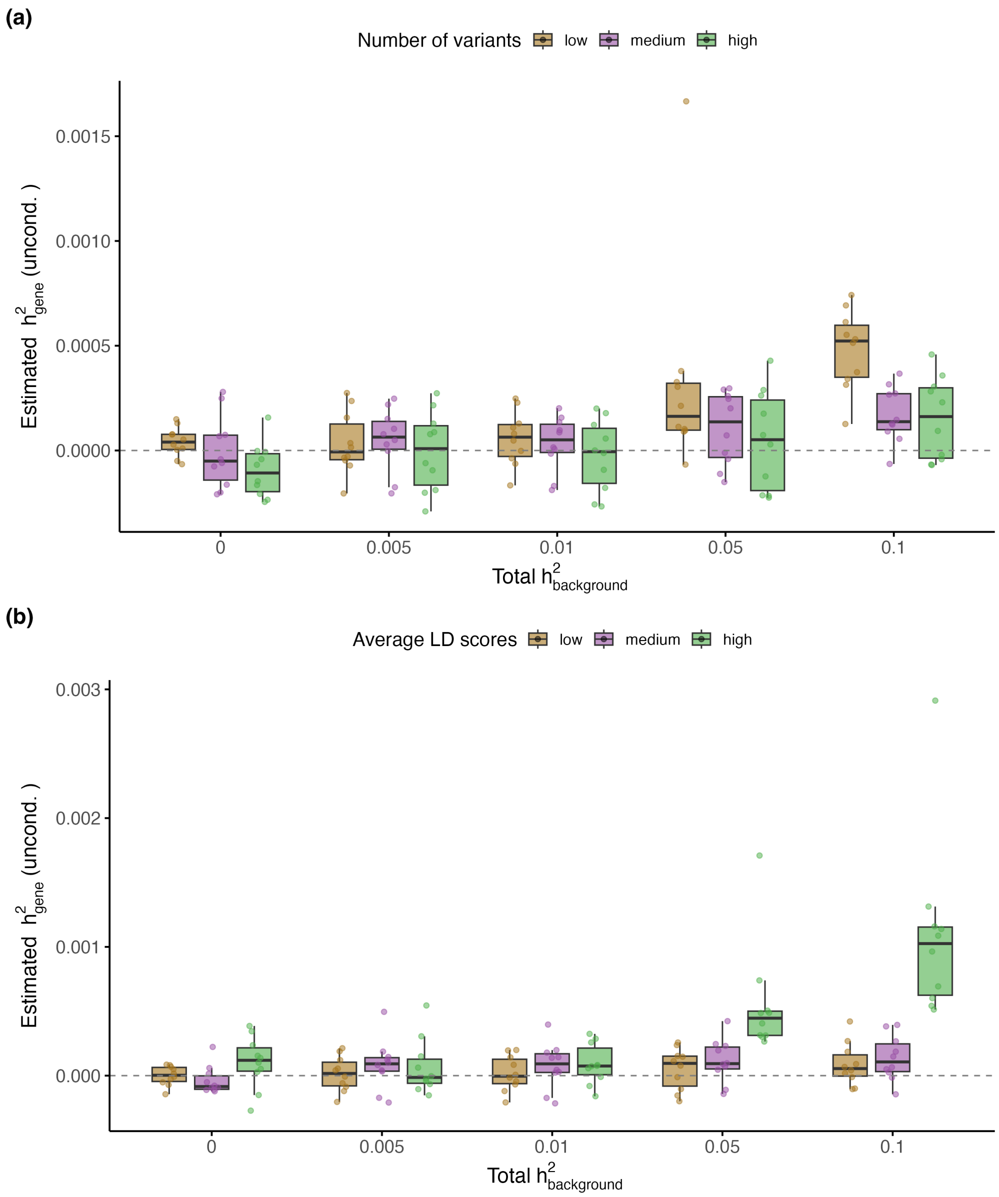


Figure S24: **LD score and gene size drive inflation in gene-level heritability estimates under the null.** **(a)** Estimated $\hat{h}^2_{\text{gene}}$ increases with background heritability ($h^2_{\text{background}}$) for genes of varying sizes, with smaller genes showing greater inflation due to reduced variant counts. **(b)** Genes with elevated LD scores exhibit more pronounced inflation under increasing $h^2_{\text{background}}$, while genes with lower LD scores remain well-calibrated. Simulations were performed under the null (no causal variants in the gene) across 10 replicates per scenario, using selected protein-coding genes on chromosome 21 representing maximum, median, and minimum values of LD score or gene size while controlling for the other parameter. FLEX was applied without LD conditioning.

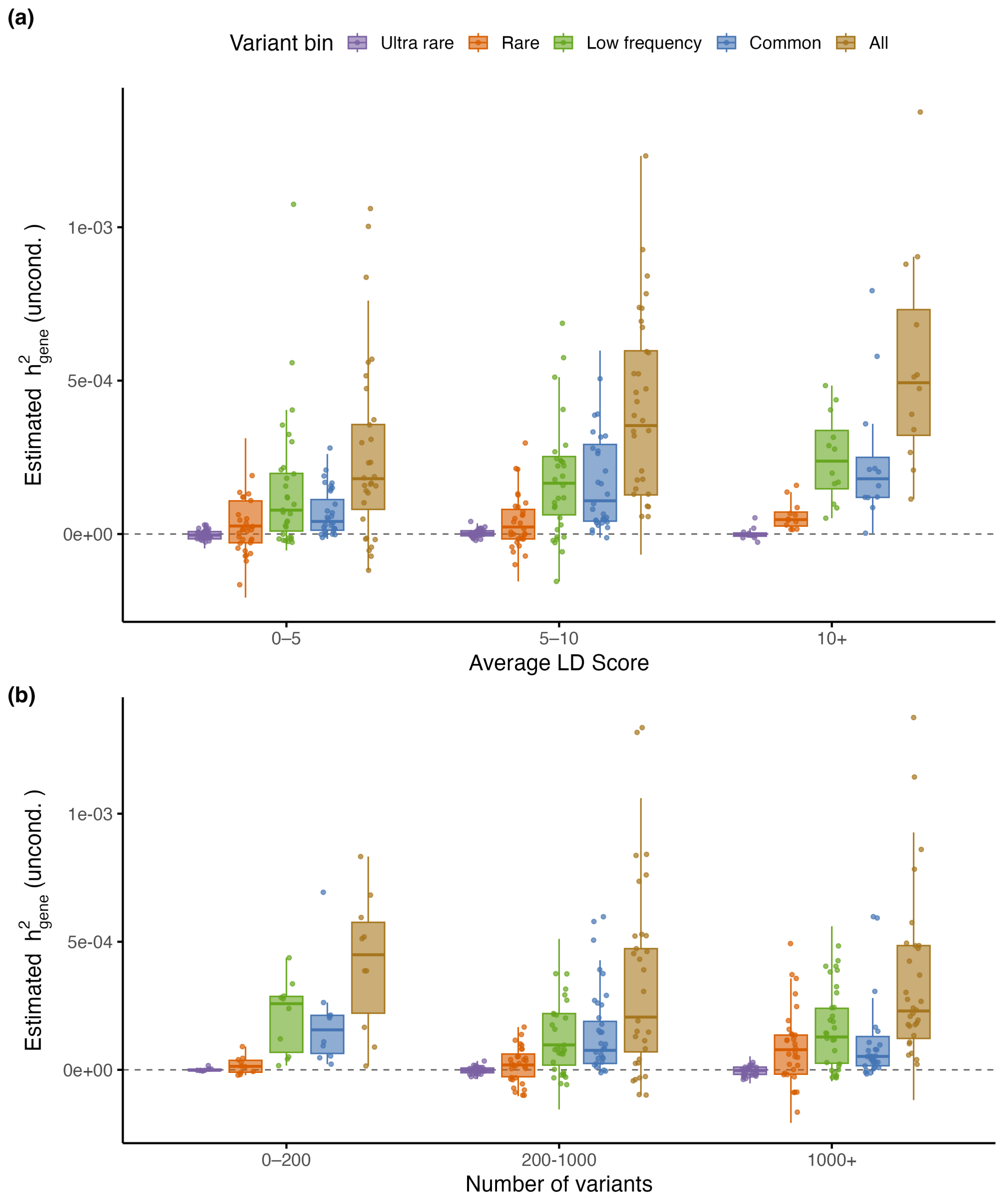


Figure S25: **Impact of LD score and gene size on heritability inflation across MAF bins. (a)** Estimated gene-level heritability ($h^2_{\text{gene}}$) stratified by average LD score and MAF bin under a background heritability of $h^2_{\text{background}} = 0.1$, with no causal variants in the gene. Inflation is observed for rare, low-frequency, and common variants in genes with high LD scores, whereas ultra-rare variants remain largely unaffected. **(b)** $h^2_{\text{gene}}$ stratified by the number of variants per gene, also under $h^2_{\text{background}} = 0.1$. Genes with fewer than 200 variants show upward bias across all MAF bins except ultra-rare, highlighting the role of gene size in LD-induced inflation. Results reflect averages over 10 replicates across 117 protein-coding genes on chromosome 21, estimated using FLEX without LD conditioning.

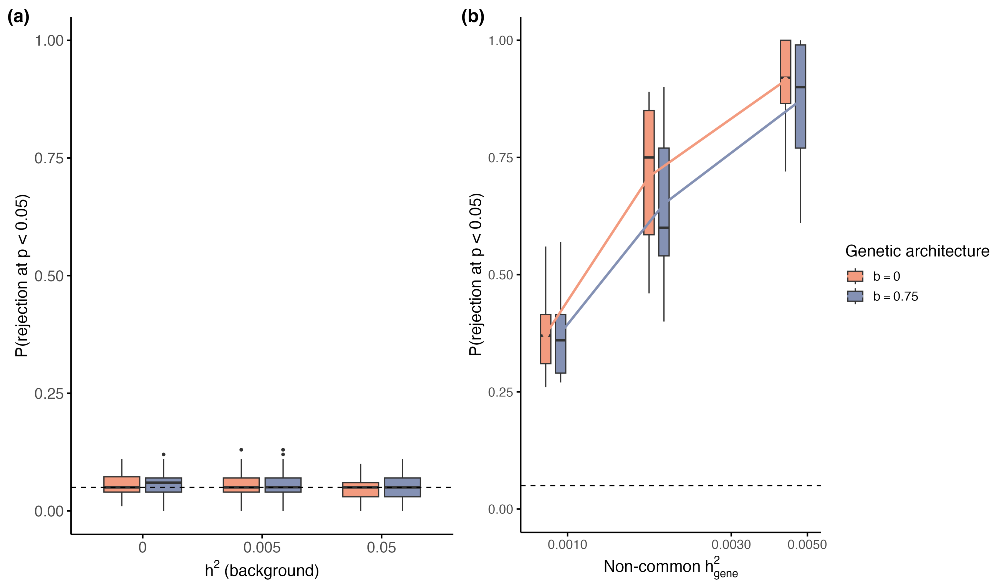


Figure S26: **Calibration and power of FLEX for binary traits.** **(a)** Calibration of $h^2_{\text{gene}}$ estimates for non-causal genes under varying $h^2_{\text{background}}$ and genetic architectures. Phenotypes were simulated using a liability threshold model and dichotomized to binary outcomes with a prevalence of 0.1. **(b)** Statistical power, defined as the proportion of causal genes detected at $p < 0.05$. Binary phenotypes were simulated with fixed chromosome-wide common heritability ($h^2_{\text{common}} = 0.01$) and varying the sum of non-common heritability of causal genes from 0.01 to 0.05, corresponding to non-common $h^2_{\text{gene}} \in [9.09 \times 10^{-4}, 4.55 \times 10^{-3}]$.
